# Identification of X-chromosomal genes that drive global X-dosage effects in mouse embryonic stem cells

**DOI:** 10.1101/2020.03.09.983544

**Authors:** Oriana Genolet, Anna A. Monaco, Ilona Dunkel, Michael Boettcher, Edda G. Schulz

## Abstract

X-chromosomal genes contribute to sex differences, in particular during early development, when both X chromosomes are active in females. Here, double X-dosage shifts female pluripotent cells towards the naive stem cell state by increasing pluripotency factor expression, inhibiting the differentiation-promoting MAP kinase (MAPK) signalling pathway and delaying differentiation. To identify the genetic basis of these sex differences, we have performed a series of CRISPR knockout screens in murine embryonic stem cells to comprehensively identify X-linked genes that cause the female pluripotency phenotype. We found multiple genes that act in concert, among which Klhl13 plays a central role. We show that this E3 ubiquitin ligase substrate adaptor protein promotes pluripotency factor expression, delays differentiation and represses MAPK target genes, and we identify putative substrates. We thus elucidate the mechanisms that drive sex-induced differences in pluripotent cells with implications for gender medicine in the context of induced pluripotent stem cell based therapies.

## Introduction

Chromosomal dosage can be altered through loss or gain of chromosomes, which, for autosomes, is generally associated with pathologies. Differential dosage of the mammalian sex chromosomes, by contrast, drives sex determination, in case of the Y, and contributes to sex differences, in case of the X chromosome (Snell and Turner 2018; Ratnu et al. 2017). The dosage imbalance for X-chromosomal genes between XX females and XY males is largely neutralized in somatic cells through X-chromosome inactivation (XCI), where one X chromosome is nearly completely silenced in each female cell (Schulz and Heard 2013). A subset of genes escape XCI and likely contribute to sex differences, for example in the context of immunity and autoimmune diseases (Schurz et al. 2019; Klein and Flanagan 2016; Wijchers et al. 2010). During early embryonic development, however, prior to the onset of XCI, the majority of X-linked genes are expressed at double the levels in female compared to male cells, resulting in substantial sex differences in cell state and developmental progression (Schulz 2017).

In many mammalian species, including mice, cows and humans, female embryos develop more slowly than their male counterparts during early development (Mittwoch 1993). Since no fetal hormones are produced at this stage, these observations have been attributed to variations in sex-chromosomal dosage, which in mice has been confirmed by the analysis of X-monosomic XO embryos (Burgoyne et al. 1995; Thornhill and Burgoyne 1993). These sex differences have been investigated at the molecular level in female mouse embryonic stem cells (mESC), which are derived from early blastocyst embryos and thus carry two active X chromosomes. Female mESCs appear to be shifted towards a more naive ground state of pluripotency, which is associated with reduced activity of the differentiation-promoting MAP kinase (MAPK) signalling pathway, increased levels of (naive) pluripotency factors and lower levels of global DNA methylation (Schulz et al. 2014; Song et al. 2019; Zvetkova et al. 2005). As a consequence, exit from the pluripotent state during differentiation is delayed in female compared to male mESCs (Schulz et al. 2014). Similar patterns have been observed in induced pluripotent stem cells (iPSCs) (Song et al. 2019). These X-dosage effects are likely mediated by X-encoded genes that modulate the stem cell state, the identity of which however remains mostly unknown. They might pose a biological checkpoint to ensure that only cells that have successfully inactivated one of their X chromosomes contribute to the differentiated adult organism.

In somatic cell types, MAPK signaling plays a key role in the regulation of cellular programs such as proliferation, but in mESCs it drives the exit from the pluripotent state, while its inhibition stabilizes the self-renewing naive ground state of pluripotency (Lanner and Rossant 2010; Ying et al. 2008). The main growth factors that stimulate MAPK signalling at these early developmental stages belong to the fibroblast growth factor (Fgf) family (Kunath et al. 2007; Stavridis et al. 2010). Upon activation of the FGF receptor (FgfR), and the subsequent membrane recruitment of the growth factor receptor-bound protein 2 (Grb2), the small GTPase Ras is activated (Kouhara et al. 1997; Lowenstein et al. 1992). Ras in turn triggers the kinase cascade of Raf, Mek and Erk. Erk then translocates to the nucleus and activates MAPK target genes, including Egr1 and Spry4 (Hodge et al. 1998; Casci et al. 1999) (Fig. S1A). Female mESCs express MAPK target genes at reduced levels compared to their male counterparts, suggesting an inhibition of the pathway (Schulz et al. 2014). To maintain homeostasis, the MAPK pathway is controlled by strong negative feedback loops on multiple levels (Lake et al. 2016). MAPK inhibition therefore often leads to a counter-intuitive rise in phosphorylation levels of pathway intermediates due to reduced negative feedback activity (Fritsche-Guenther et al. 2011; Sturm et al. 2010). Female mESCs, where the MAPK pathway is inhibited, thus exhibit increased Mek phosphorylation compared to male cells, suggesting inhibition of the pathway downstream of Mek (Schulz et al. 2014; Choi et al. 2017).

MAPK signaling and pluripotency are tightly coupled, as the inhibition of this pathway blocks differentiation, leads to an increased expression of naive pluripotency markers and DNA hypomethylation, a hallmark of the naive pluripotent state (Ficz et al. 2013; Kunath et al. 2007; Silva et al. 2009; Niwa et al. 2009). Reduced MAPK signalling in female mESCs thus results in increased levels of naive pluripotency factors such as Nanog and Prdm14, and global DNA hypomethylation (Zvetkova et al. 2005; Ooi et al. 2010; Ficz et al. 2013; Leitch et al. 2013; Habibi et al. 2013; Hackett et al. 2013; Milagre et al. 2017; Pasque et al. 2018; Schulz et al. 2014).

Although X-chromosomal dosage exhibits global effects on signalling and gene expression, central X-encoded genes that mediate these phenotypes remain to be uncovered (Schulz 2017). The X-linked Erk-phosphatase Dusp9 has been shown to underlie sex differences in DNA methylation, since a heterozygous mutation resulted in DNA hypermethylation as observed in male cells (Caunt and Keyse 2013; Choi et al. 2017). However, pluripotency factor expression and differentiation has been reported to be unaffected in such mutant cells (Song et al. 2019). Moreover, a series of other X-linked genes, including the transcription factors Zic3 and Tfe3 have been investigated, but their heterozygous deletion in female cells had no detectable effect (Song et al. 2019). Taken together, the genetic determinants that drive sex differences in mESCs remain incompletely understood.

We have performed a series of complementary CRISPR screens to identify X-linked genes that modulate MAPK signalling, pluripotency and differentiation and found several genes that contribute to these phenotypes. We show that the E3 ubiquitin ligase adaptor protein Klhl13 promotes pluripotency factor expression, while inhibiting MAPK target gene expression and differentiation. Female mESCs carrying heterozygous mutations of Klhl13 and the known X-linked MAPK inhibitor Dusp9 qualitatively recapitulate all aspects of the male pluripotency phenotype. We have thus identified the main drivers of X-dosage-dependent sex differences in mESCs and disentangled their relative contributions. Our approach can serve as a blueprint to investigate dosage effects of other chromosomes, such as those underlying trisomy 21, and our results will be important for development of gender-sensitive iPSC-based therapies.

## Results

### Pooled CRISPR knockout screen reveals X-chromosomal MAPK regulators

The X chromosome encodes ∼1000 genes, any of which could potentially mediate the sex differences observed in murine pluripotent stem cells with respect to pluripotency factor expression, MAPK pathway activity and differentiation efficiency (Schulz et al. 2014; Song et al. 2019). Since MAPK signalling represses pluripotency factors and promotes differentiation (Kunath et al. 2007; Silva et al. 2009; Niwa et al. 2009), we hypothesized that an X-linked MAPK inhibitor might underlie the female pluripotency phenotype (Schulz 2017; Schulz et al. 2014). To comprehensively identify X-encoded MAPK inhibitors, we performed a chromosome-wide pooled CRISPR knockout screen (Fig. 1A). Through transduction of Cas9-expressing mESCs with an X-chromosomal sgRNA expression library, a pool of cells was generated with maximally one gene mutated per cell. Subsequent enrichment of cells with increased MAPK pathway activity and sequencing of their associated sgRNAs allowed identification of genes acting as MAPK inhibitors that, when deleted, increased MAPK signalling.

**Figure 1.**
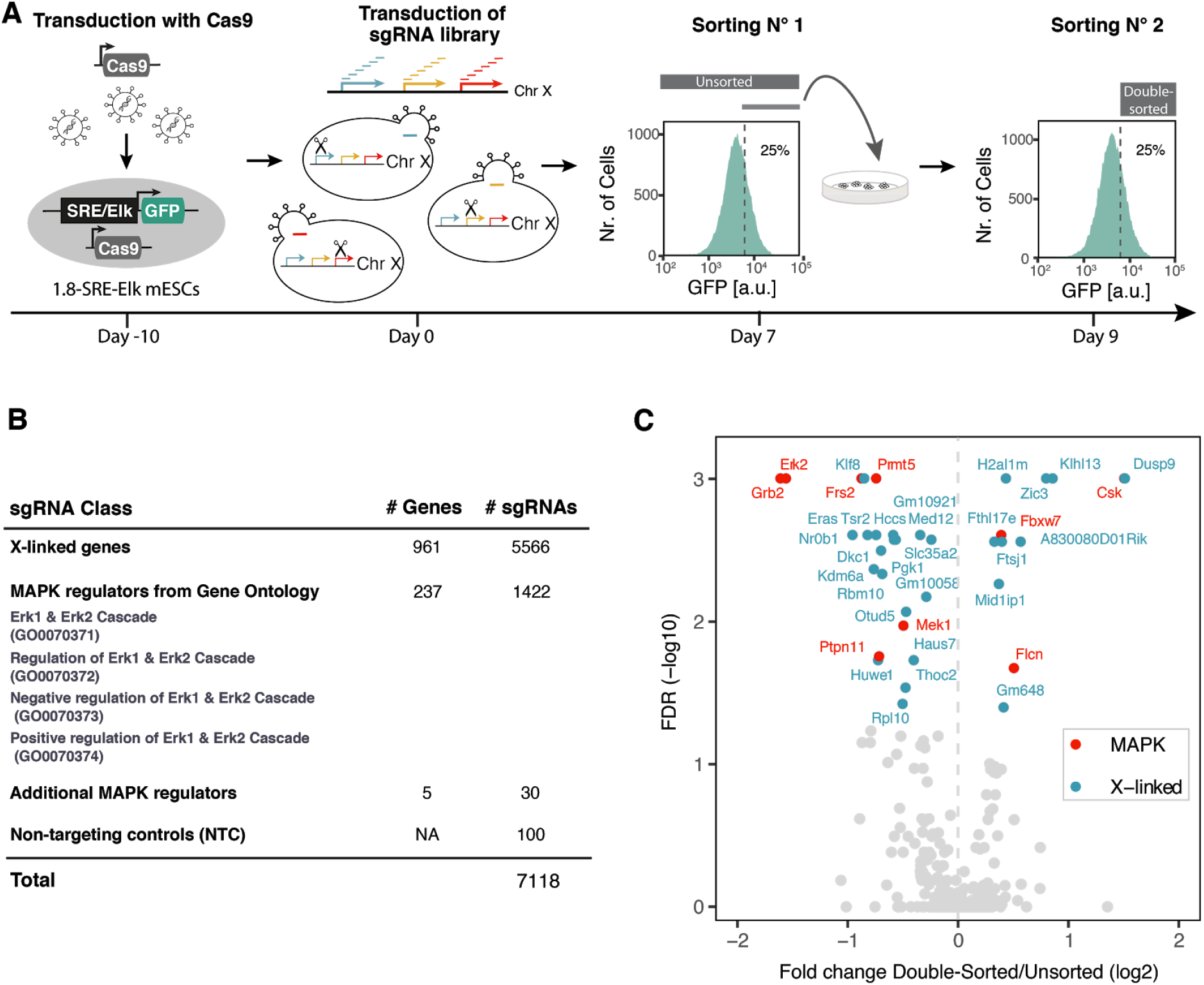
Identification of X-chromosomal MAPK regulators through a pooled CRISPR knockout screen. **(A)** Schematic depiction of the screen workflow: A female mESC line carrying a stably integrated fluorescent MAPK reporter, where expression of GFP is controlled by an SRE-Elk responsive promoter, was transduced with a construct expressing the Cas9 endonuclease. Cells were further transduced with a custom sgRNA library targeting the majority of X-chromosomal genes. GFP-high cells were sorted by flow cytometry, cultured for an additional 2 days and sorted again (double-sorted). The sgRNA cassette was amplified from genomic DNA and sgRNA abundance in the unsorted and double-sorted populations was determined by deep sequencing. The screen was performed in three independent replicates. **(B)** Composition of the GeCKOx sgRNA library, targeting X-linked genes and positive control genes known to regulate the MAPK pathway, with 6 sgRNAs per gene. As negative controls, non-targeting sgRNAs were included in the library. **(C)** Volcano plot of the screen results, where screen hits (FDR<0.05, MAGeCK) are labelled in red (positive controls) or blue (X-linked genes).

To be able to enrich cells with high MAPK activity through fluorescence-activated cell sorting (FACS), we generated a female mESC line (1.8-SRE-Elk), where expression of GFP was driven by a synthetic MAPK-sensitive SRE-Elk promoter (containing binding sites for the transcription factors Elk1 and Srf, which are activated downstream of the MAPK pathway) (Fig. 1A). Reporter functionality was confirmed by treatment with an inhibitor of Mek, which resulted in the expected decrease in GFP fluorescence (Fig. S1B). To focus the screen on X-linked genes, we generated a custom sgRNA library (GeCKOx) containing a subset of sequences of the genome-wide GeCKO library (Shalem et al. 2014), targeting 961 X-chromosomal genes with 6 sgRNAs per gene, where possible. As controls, 237 genes implicated in MAPK pathway regulation according to gene ontology (GO) annotation and 100 non-targeting controls (NTC) were included in the library (Fig. 1B, Supp. Table S1). Sequencing of the sgRNA library confirmed an even representation (Fig. S1C).

To investigate the female pluripotency phenotype, cells were generally grown in classical ESC culture conditions, containing Serum and LIF, if not stated otherwise. For the screen, 1.8-SRE-Elk mESCs were first transduced with a lentiviral vector expressing the Cas9 endonuclease, followed by blasticidin selection, sgRNA library transduction and puromycin selection. After expansion for 7 days (5 days under selection), cells with high reporter activity were FACS-sorted, replated and cultured for two additional days, before being sorted once again (Fig. 1A). We reasoned that such a double-sorting strategy would increase sensitivity of the screen. The sgRNA cassette was amplified from genomic DNA of all double-sorted (day 9) and unsorted (day 7) cell populations and sgRNA abundance in each sample was quantified by Illumina sequencing. SgRNA counts in all libraries were highly correlated and NTCs were neither enriched nor depleted in the sorted fractions, suggesting that sufficient coverage was maintained at all steps of the screen (Fig. S1D-F).

Several core MAPK pathway components were significantly depleted in the GFP-high population (Erk2, Grb2, Frs2, Mek1 and Ptpn11), while Csk, a MAPK inhibitor (Okada 2012), was enriched, showing that our screening setup could recover positive controls (Fig. 1C, red, Supp. Table S1). Among the X-linked genes, 9 were significantly enriched and 18 were depleted in the sorted population (FDR < 0.05, MAGeCK, Fig. 1C, blue). Dusp9, Klhl13 and Zic3 were the top-scoring MAPK inhibitors and Klf8, Nr0b1 and Eras were the strongest activators (Fig. 1C).

In principle, enrichment in the double-sorted fraction at day 9 compared to the unsorted cells at day 7 could also be due to faster proliferation between the two sampling points. To identify genes that affect proliferation or viability, we compared sgRNA frequency in the cloned library and the unsorted cells at day 7 (Fig. S1G, Supp. Table S1). Among the identified X-linked MAPK inhibitors, only H2al1m seemed to affect mESC proliferation positively, which would however lead to a decrease and not an enrichment, in sgRNA abundance between day 7 and day 9. In summary, we found a series of X-encoded inhibitors of the MAPK pathway, which might potentially drive the X-dosage-dependent pluripotency phenotype.

### Secondary screens identify X-linked regulators of pluripotency factors, differentiation kinetics and Mek phosphorylation

Having identified a set of putative X-linked MAPK pathway regulators, we further investigated their function in a series of complementary CRISPR screens. Specifically, we tested whether the identified candidate genes affected pluripotency factor expression, differentiation dynamics and phosphorylation of Mek in a manner that would phenocopy the male pluripotency phenotype. For this purpose, a sub-library of the GeCKOx sgRNA library (GeCKOxs) was generated, targeting the 50 most enriched and depleted X-linked genes, together with the 10 most enriched and depleted MAPK controls from the primary screen (Fig. 2A, Fig. S2A, Supp. Table S1). For each gene, the three most effective sgRNAs were selected. In addition, sgRNAs targeting 10 pluripotency regulators were included as further controls (Sox2, Tbx3, Tcf3, Fgf2, Stat3, Esrrb, Tfcp2l1, Klf2, Nanog and Oct4).

**Figure 2.**
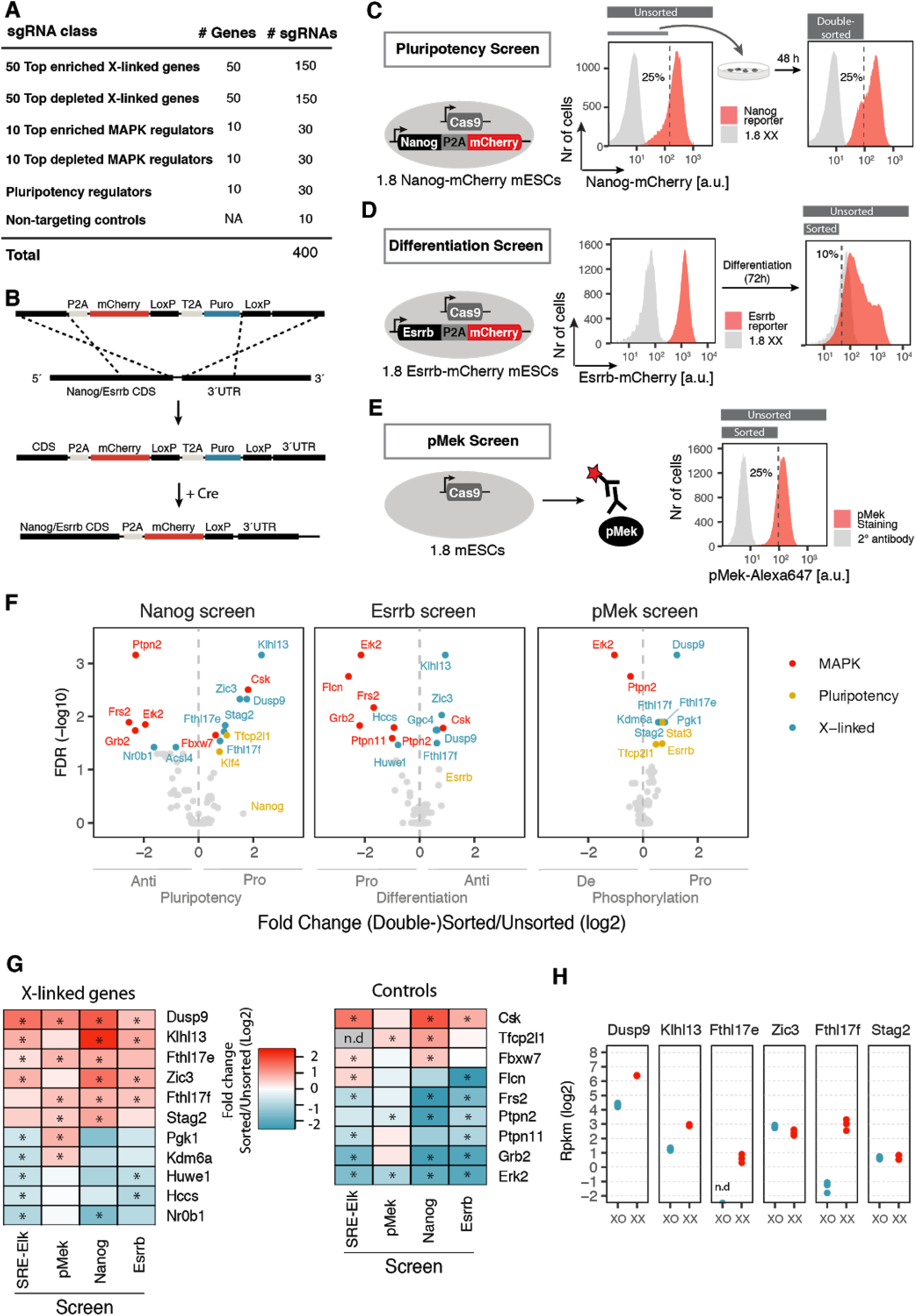
Secondary CRISPR screens profiling pluripotency factor expression, differentiation kinetics and Mek phosphorylation. **(A)** Composition of the GeCKOxs sgRNA library, targeting hits from the primary MAPK screen and positive control genes with 3 sgRNAs per gene. **(B)** Schematic representation of the C-terminal tagging of the *Nanog* and *Esrrb* genes with the mCherry fluorescent protein through CRISPR/Cas9-mediated homologous recombination and subsequent Cre-mediated excision of the puromycin resistance cassette. Nanog/Esrrb and mCherry are linked through a P2A self-cleaving peptide. **(C-E)** Schematic depiction of the three secondary screens to profile effects on pluripotency factor expression (C), differentiation (D) and Mek phosphorylation (E). Female mESCs, carrying mCherry-tagged *Esrrb/Nanog* loci, as indicated, expressing the Cas9 endonuclease, were transduced with the sgRNA library in (A). **(C)** In the Nanog screen, the 25% cells with the weakest mCherry fluorescence were enriched in two consecutive sorts (day 7 and day 9 after transduction). **(D)** For the Esrrb screen, cells were differentiated via LIF withdrawal for 3 days and the 10% cells with the lowest mCherry fluorescence were FACS sorted. **(E)** In the pMek screen, cells were stained intracellularly with a pMek-specific antibody and the 25% cells with the lowest signal were sorted. Three replicates were generated for the Esrrb and pMek reporter screens and two for the Nanog screen. **(F)** Volcano plot of the most enriched and depleted genes in the Nanog, Esrrb and pMek screens. Genes with an FDR<0.05 are highlighted as indicated. **(G)** Heatmap summarizing the results from all 4 screens. Enrichment of all X-linked (left) and control genes (right) that were significantly enriched or depleted in at least 2 screens is shown. *FDR<0.05 (MAGeCK), n.d non-determined **(H)** Expression levels for a subset of X-linked genes shown in (G) in 1.8XX and 1.8XO mESCs assessed by RNA-sequencing.

To assess effects on pluripotency factor expression, we decided to assay for Nanog levels, which are consistently higher in female compared to male mESCs (Schulz et al. 2014; Choi et al. 2017; Song et al. 2019). As a readout for differentiation efficiency, we monitored Esrrb, a naive pluripotency marker, which is down-regulated with faster dynamics in cells with only one X chromosome (Schulz et al. 2014). We generated two transgenic mESC lines, where the endogenous *Nanog* and *Esrrb* genes, respectively, were tagged C-terminally with the fluorescent protein mCherry (Fig. 2B, Fig. S2B-C). Both reporters were down-regulated upon differentiation, suggesting that they indeed mirrored expression of Nanog and Esrrb, respectively (Fig. S2D). In the pluripotency screen we aimed at identifying Nanog activators, which when knocked-out would reduce Nanog levels, and therefore sorted cells with low Nanog expression (Fig. 2C). Similarly, the differentiation screen aimed at identifying genes that would, when deleted, induce a more rapid down-regulation of Esrrb. We therefore sorted Esrrb-low cells after 3 days of differentiation (Fig. 2D). For the Nanog screen, a double-sorting strategy similar to the primary MAPK screen was used, while only a single sorting step was performed for the differentiation screen, where a transient phenotype was analyzed (Fig. 2C-D). In a third secondary screen we aimed to test whether deletion of the candidate genes would result in decreased phosphorylation of Mek as observed in cells with one X chromosome (Schulz et al. 2014; Choi et al. 2017). To this end, we performed an intra-cellular staining with a pMek-specific antibody and sorted cells with a low pMek signal (Fig. 2E). Staining specificity was confirmed by a higher pMek signal in XX compared to XO cells, together with an increase in pMek levels upon Meki treatment in the latter (Fig. S2E). Since the staining required cell fixation only a single sorting step was possible. Sufficient sgRNA library representation was maintained throughout all steps of the screens (Fig. S2F). NTCs were neither enriched nor depleted in the pluripotency and differentiation screens, but seemed slightly but significantly depleted in the pMek-screen (Fig. S2G). SgRNAs targeting the screen hits however exhibited a much stronger effect (Fig. S2G). Among the known MAPK regulators, the pathway components Erk2, Grb2 and Frs2 were identified as anti-pluripotency and pro-differentiation factors and the negative MAPK regulator Csk showed the opposite behavior (Fig. 2F, red, Supp. Table S1). Erk2 also scored as the strongest negative regulator of Mek phosphorylation due to strong Erk-mediated negative feedback regulation (Sturm et al. 2010; Fritsche-Guenther et al. 2011). Ptpn2, a known negative regulator of MAPK signaling (van Vliet et al. 2005; Mattila et al. 2005), was surprisingly identified as an anti-pluripotency and pro-differentiation factor, potentially due to its previously reported inhibitory effect on Jak/STAT signaling, a pro-pluripotency pathway (Yamamoto et al. 2002). Moreover, Folliculin (Flcn) was identified as a strong pro-differentiation factor in agreement with its previously reported central role in early differentiation (Betschinger et al. 2013). Finally, also the pluripotency factors Tfcp2l1 and Klf4 were identified as Nanog activators as expected (Martello et al. 2013; Hall et al. 2009; Niwa et al. 2009). Nanog and Esrrb themselves were enriched 3.1 (FDR=0.67) and 1.6-fold (FDR=0.2), respectively. The low statistical power to detect Nanog enrichment can be attributed to the fact that sgRNAs targeting Nanog become depleted over time, because they affect proliferation (Fig. S2H, Supp. Table S1). Interestingly, the pluripotency regulators Stat3, Esrrb and Tfcp2l1 scored as positive regulators of Mek phosphorylation, potentially in part due to crosstalk from the Jak/Stat to the MAPK signaling pathway (Cacalano et al. 2001). In summary, all three secondary screens recovered known regulators supporting the validity of the approaches.

In all three secondary screens 5-6 X-linked genes were enriched in the (double-)sorted populations, while only maximally 2 were depleted (Fig. 2F, blue, Fig. 2G, Supp. Table S1). The only gene that significantly affected all 4 phenotypes (including the SRE-Elk screen, Fig. 1) was Dusp9, a known MAPK inhibitor that de-phosphorylates Erk and has previously been implicated in sex differences in ES cells (Choi et al. 2017; Caunt and Keyse 2013; Song et al. 2019). In addition, Klhl13, two members of the Fthl17 cluster, Fthl17e and Fthl17f, Zic3 and Stag2 significantly affected 2-3 phenotypes and generally showed the expected trend in all screens (Fig. 2G, Fig. S2I). Taken together, we have identified 6 genes that might contribute to the sex differences observed in mESCs, none which, apart from Dusp9, has previously been implicated in mediating sex differences. Klhl13 encodes a substrate adaptor protein for the Cullin3 E3 ubiquitin-protein ligase complex with no known role in pluripotency or MAPK signalling regulation (Sumara et al. 2007). The Fthl17 gene cluster encodes ferritin-like proteins with unknown functions that are partially nuclear and lack ferroxidase activity (Ruzzenenti et al. 2015). Zic3 is a transcription factor implicated in pluripotency and early differentiation, whereas Stag2 regulates chromatin conformation and has also been shown to be involved in the maintenance of the pluripotent state in mESCs (Lim et al. 2007, 2010; Kagey et al. 2010; Yang et al. 2019). Among these candidates the strongest effects were observed for Dusp9 and Klhl13.

### Klhl13 and Dusp9 exhibit higher levels in females *in vitro* and *in vivo*

To further characterize the six identified putative mediators of the female pluripotency phenotype, we compared their expression pattern between cells with one and two X chromosomes, both *in vitro* and *in vivo*. We generated RNA-sequencing data of the female mESC line used in all screens (1.8XX) and a subclone of that line with only one X chromosome (1.8XO). Although X-linked genes showed in general the expected 2-fold higher expression in XX compared to XO cells (Fig. S3A), two genes, Zic3 and Stag2, were expressed at similar levels in the two cell lines (mean fold change 0.8 and 1.1 respectively), potentially due to gene-specific dosage-compensation mechanisms (Fig. 2H, Supp. Table S2). Dusp9 and Klhl13 were expressed at 4.2- and 3.2-fold higher levels in XX compared to XO cells, respectively, and the two members of the Fthl17 cluster were essentially not expressed in the XO line (Fig. 2H). The strong expression difference for Fthl17e and Fthl17f can be explained by the fact that the cluster is maternally imprinted, such that it is only expressed from the paternal X chromosome, which is present only in female embryos and was probably also lost in the XO clone (Kobayashi et al. 2010).

To assess expression patterns in mouse embryos *in vivo* we analyzed epiblast cells in published single-cell RNA sequencing data collected between embryonic days E4.5 and E6.5 (Fig. S3B-G) (Argelaguet et al. 2019). Reactivation of the paternal X chromosome, which is silenced early in development in an imprinted form of XCI, is initiated around E4.5, completed at E5.5 and followed by random XCI around E6.5 (Mohammed et al. 2017; Cheng et al. 2019). X-chromosomal expression was thus 1.6- and 1.4-fold higher in female compared to male cells at E4.5 and E5.5, respectively, with the difference being largely neutralized by E6.5 (Fig. S3B). In contrast to mESCs, where both X chromosomes are active in the naive pluripotent state, *in vivo* naive pluripotency factors are primarily expressed prior to X reactivation around E3.5 and are mostly downregulated at E4.5 (Mohammed et al. 2017). As a consequence, most naive markers were not well detected in the data set we analyzed and a combined analysis of 9 naive factors revealed only a slight trend towards higher expression in female cells at E4.5 (Fig. S3C-D) Analysis of a group of 9 markers of the primed pluripotent state, by contrast, showed a clear trend towards higher expression in all three time points (Fig. S3E-F), suggesting that differentiation of female cells with a double X-dosage is also delayed in embryos *in vivo*. Analysis of the six identified putative candidate genes revealed a trend towards higher expression in female cells at E5.5 for all factors, which was statistically significant (p<0.05, Wilcoxan ranksum test) for Dusp9, Fthl17e, Fthl17f and Stag2 (Fig. S3G). In summary, all six factors were expressed at higher levels in female compared to male cells *in vivo*, but only four of them (Dusp9, Klhl13, Fthl17e/f) showed the same trend in the 1.8XX/XO cell lines *in vitro*. Since the 1.8XX/XO lines show a strong X-dosage dependent phenotype (Schulz et al. 2014), we concluded that the four differentially expressed factors would be the best candidates for mediating X-dosage effects on pluripotency and differentiation and decided to further validate Dusp9 and Klhl13, which appeared to induce the strongest phenotypes.

### Over-expression of Klhl13 and Dusp9 leads to an enhanced pluripotency state and slower differentiation kinetics in male mESCs

If Dusp9 and Klhl13 would indeed mediate the sex differences observed in mESCs, their over-expression in male cells should lead to a female-like pluripotency phenotype, while their heterozygous deletion should shift female cells towards a male-like phenotype. In order to over-express Klhl13 and Dusp9 from their endogenous loci in male mESCs we implemented the CRISPR activation (CRISPRa) system. We made use of an E14 mESC line carrying the components of the CRISPRa SunTag system under control of a doxycycline-inducible promoter, which allows recruitment of multiple VP64 activation domains through a single sgRNA (Fig. 3A) (Tanenbaum et al. 2014; Heurtier et al. 2019).

**Figure 3.**
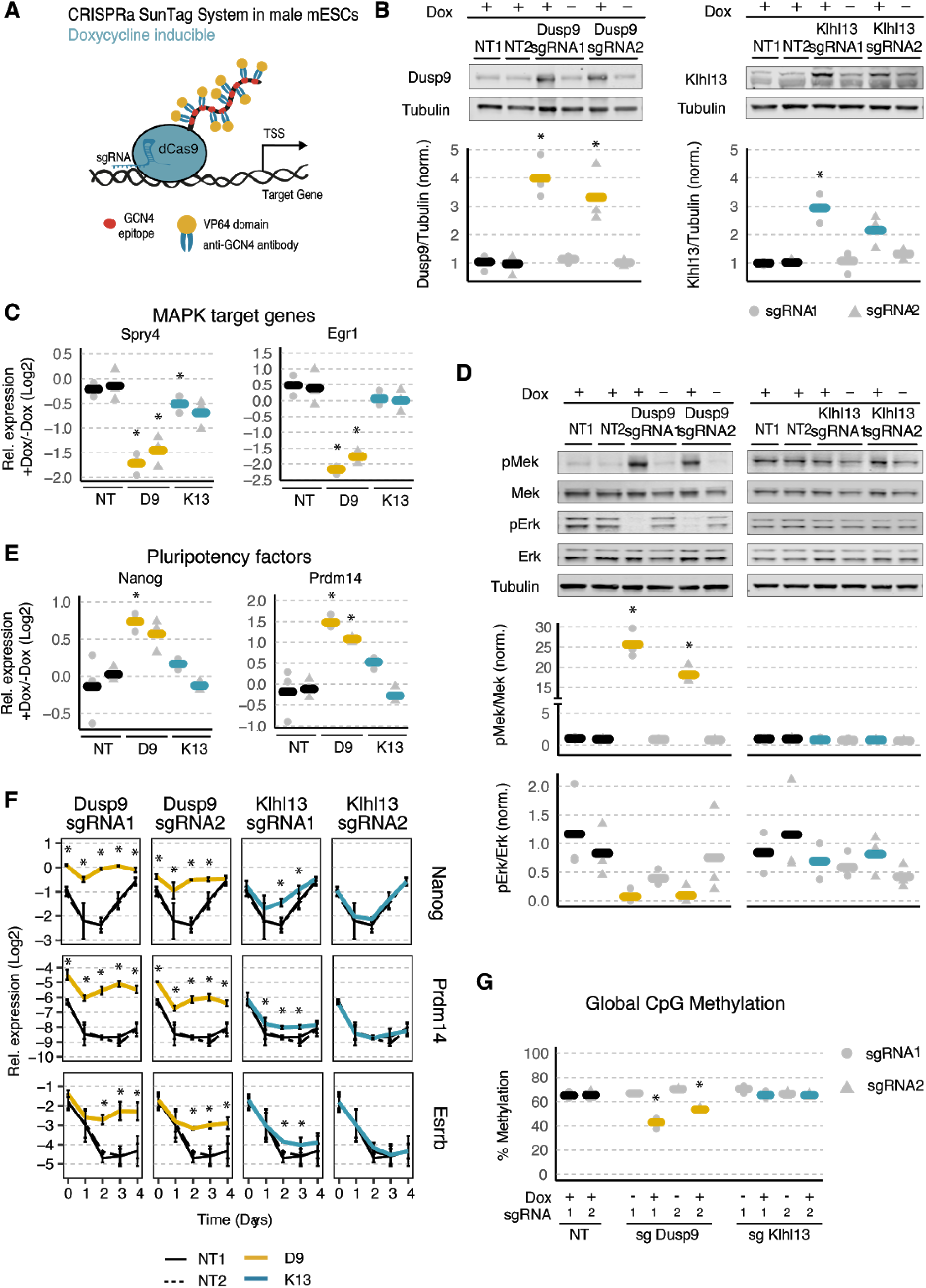
Over-expression of Klhl13 and Dusp9 in male mESCs leads to an enhanced pluripotency state and slower differentiation kinetics. **(A)** Schematic representation of the dCas9-SunTag system used for gene activation. **(B-E)** To over-express Dusp9 (yellow) and Klhl13 (blue), male E14 mESCs, stably expressing the doxycycline-inducible SunTag system, were either transduced with one of two different sgRNAs targeting the respective promoter regions or with non-targeting control (NT) sgRNAs and were treated for 3 days with 1 µg/ml doxycycline as indicated. Protein levels of Dusp9 (left) and Klhl13 (right) were quantified via immunoblotting (B), expression levels of MAPK target genes Spry4 and Egr1 (C) and of naive pluripotency factors Nanog and Prdm14 (E) were assessed by qPCR and phosphorylation of Mek and Erk was quantified by immunoblotting (D). The immunoblot signals were normalized to Tubulin (B) or to total Mek/Erk (D) and to the mean of two doxycycline-treated non-targeting control sgRNAs. qPCR measurements were normalized to two housekeeping genes and to the respective untreated control (-Dox). Dots and triangles depict individual measurements of the two different sgRNAs, and thick bars show the mean of three biological replicates. **(F)** Dusp9- and Klhl13 over-expressing mESCs were treated with 1 µg/ml doxycycline 24 h before differentiation via LIF withdrawal for 4 days and expression levels of pluripotency factors were measured by qPCR at different time points as indicated. Mean and standard deviation across 3 biological replicates is shown. **(G)** Global CpG methylation levels in cell lines over-expressing Dusp9 and Klhl13 via doxycycline treatment for 3 passages were assessed via pyrosequencing-based luminometric DNA methylation assay (LUMA). * p < 0.05 in a two-tailed paired Student’s t-test comparing the Dusp9/Klhl13 over-expressing samples and the non-targeting controls (mean of sgRNA1 and sgRNA2).

The SunTag system was recruited to either the Dusp9 or Klhl13 promoters using two different sgRNAs per gene and one sgRNA per cell line, leading to a 4- and 3.3-fold over-expression of Dusp9 protein and to a 2.9- and 2.1-fold induction of Klhl13 protein, respectively (Fig. 3B, Fig. S4A). We then characterized these cell lines with respect to pluripotency factor expression, differentiation dynamics, MAPK pathway activity and global DNA methylation levels, all of which are affected by X-chromosomal dosage in mESCs. To assess MAPK pathway activity we measured expression levels of Spry4 and Egr1 (Hodge et al. 1998; Casci et al. 1999), two well known Erk target genes, by qPCR (Fig. 3C). Both MAPK target genes were strongly down-regulated upon Dusp9 over-expression (2.7/5.5-fold), as expected for an Erk phosphatase, while their expression was only slightly, albeit mostly not significantly reduced upon Klhl13 over-expression. When assessing phosphorylation levels of MAPK pathway intermediates, we found that Dusp9 over-expression reduced pErk levels 12-fold, again as expected for an Erk phosphatase, but increased pMek 22-fold, most possibly due to reduced negative feedback inhibition (Fig. 3D). Over-expression of Klhl13 by contrast had no significant effect on either Erk or Mek phosphorylation. A previous study had reported the opposite effect of Dusp9 over-expression on Erk phosphorylation, potentially due to the requirement for trypsinization when analyzing feeder-dependent mESC lines, which were used in that study (Choi et al. 2017) (Fig. S4B).

Taken together, these results confirm that Dusp9 is a strong inhibitor of MAPK pathway activity, while Klhl13 might slightly inhibit MAPK target gene expression, but does not affect pathway intermediates, which is in accordance with our screening results (Fig. 2G).

We next assessed how over-expression of Klhl13 and Dusp9 would affect pluripotency factor expression and differentiation dynamics. To this end, we quantified the pluripotency factors Nanog and Prdm14, which have been reported to be expressed at 2-4-fold higher levels in PSCs with two X chromosomes compared to those with one (Choi et al. 2017; Song et al. 2019; Schulz et al. 2014). Over-expression of Dusp9 in male mESCs resulted in a nearly comparable increase of Nanog (1.7-1.5-fold) and Prdm14 (3-2.3-fold) levels. Upon Klhl13 over-expression by contrast, only Prdm14 was increased (1.6-fold) and only by the stronger sgRNA (Fig. 3E). A very similar trend was observed with regard to differentiation dynamics, where Dusp9 over-expression essentially blocked down-regulation of naive pluripotency markers, Nanog, Prdm14 and Esrrb, while for Klhl13 only the stronger sgRNA had a mild effect on differentiation dynamics (Fig. 3F). Over-expression of Dusp9 in male cells thus seemed to induce a strong shift towards the naive pluripotent state similar to female cells, while Klhl13 over-expression resulted in only a minor shift.

Since Dusp9 has been suggested to be responsible for the reduction of global CpG methylation levels typically observed in female mESCs (20-30% compared to 60-80% in male mESCs) (Choi et al. 2017; Zvetkova et al. 2005; Habibi et al. 2013), we analzed how over-expression of Dusp9 and Klhl13 affected global DNA methylation through the pyrosequencing-based luminometric DNA methylation assay (LUMA, Fig. 3G). Upon Dusp9 over-expression, global DNA methylation levels were reduced from ∼60% in NTC-transduced control cells to 53% and 42%, but were unaffected by Klhl13 over-expression. Our results confirm a previously described effect of Dusp9 on global DNA methylation (Choi et al. 2017).

Overall we observe a stronger induction of a naive-like state in Dusp9-compared to Klhl13-overexpressing cells. It is important to note, however, that over-expression is less efficient for Klhl13 than for Dusp9 and that the observed effects seem to be strongly dose dependent. The fact that small, but significant effects are observed also for Klhl13 with the stronger sgRNA (which increases Klhl13 expression to levels similar albeit slightly lower to those in females) suggests that also Klhl13 might contribute to sex differences with respect to pluripotency factor expression, differentiation and MAPK target gene expression. To test this we further investigated the role of both genes in female mESCs.

### Mutation of one copy of *Klhl13* and *Dusp9* in female mESCs induces the male pluripotency state

If increased expression of Klhl13 and Dusp9 in female compared to male cells is indeed what drives sex differences in mESCs, their deletion on one X chromosome in female ESCs should induce the male phenotype. We therefore generated both heterozygous (HET) and homozygous (HOM) mutant mESC lines for Klhl13 (K13) and Dusp9 (D9) and a heterozygous double-mutant line (D9K13). For Klhl13, a 5kb region spanning the promoter was deleted using CRISPR-Cas9, whereas for Dusp9, where attempts to create a promoter deletion were unsuccessful, frameshift mutations were introduced through an sgRNA targeting the start of the coding sequence (CDS) (Fig. 4A, Fig. S4C-D). Two clones were analyzed for each genotype throughout all experiments except for differentiation dynamics. Loss of Klhl13 transcription in the respective mutants was confirmed by nascent RNA FISH (Fig. S4E) and all generated clones were karyotyped via double digest genotyping-by-sequencing (Fig. S4F) (Elshire et al. 2011). Dusp9 protein levels were reduced ∼1.8-fold in the respective HET mutants, which is less than the 3.5-fold reduction observed when comparing XX to XO cells, suggesting that Dups9 levels are modulated by other X-linked genes (Fig. 4B). In HET lines with a Klhl13 mutation the Klhl13 protein was reduced ∼2.7-fold (Figure 4B). In all cell lines we then analyzed MAPK signalling, pluripotency factor expression, differentiation and global DNA methylation. To assess whether MAPK pathway activity was affected in the mutant cell lines, we again quantified expression of the MAPK target genes Egr1 and Spry4 (Fig. 4C, Fig. S5A). Both were expressed at higher levels in all mutant lines compared to the XX control clones, suggesting that the MAPK pathway inhibition was at least partially lifted. Among the HET mutant lines, D9 showed the weakest effect, followed by K13 and D9K13, with the double mutant reaching similar expression levels as found in the XO control cells (3.4/6.5-fold for Spry4/Egr1 in D9K13-HET vs 2.7/10.2 in XO). To get a more global picture of signalling activity, we analyzed a larger set of MAPK target genes using RNA-sequencing (Fig. 4D, Fig. S5B, Supp. Table S2). In agreement with the qPCR results, we found MAPK target genes were significantly increased in K13-HET cells, and further elevated in D9K13 double mutants. We also assessed signatures of two other signalling pathways, Akt and Gsk3, implicated in pluripotency and differentiation, for which differential activity has been found in male and female mESC (Schulz et al. 2014); (Watanabe et al. 2006; Wray et al. 2011). Again the heterozygous D9K13 mutant cells showed the strongest effects on Akt and Gsk3 target genes (Fig. 4D, Fig. S5B). It is important to note, however, that for none of the pathways we analyzed, target gene expression reached the levels found in XO control clones, suggesting that additional genes, other than Dusp9 and Klhl13, are involved in their regulation.

**Figure 4.**
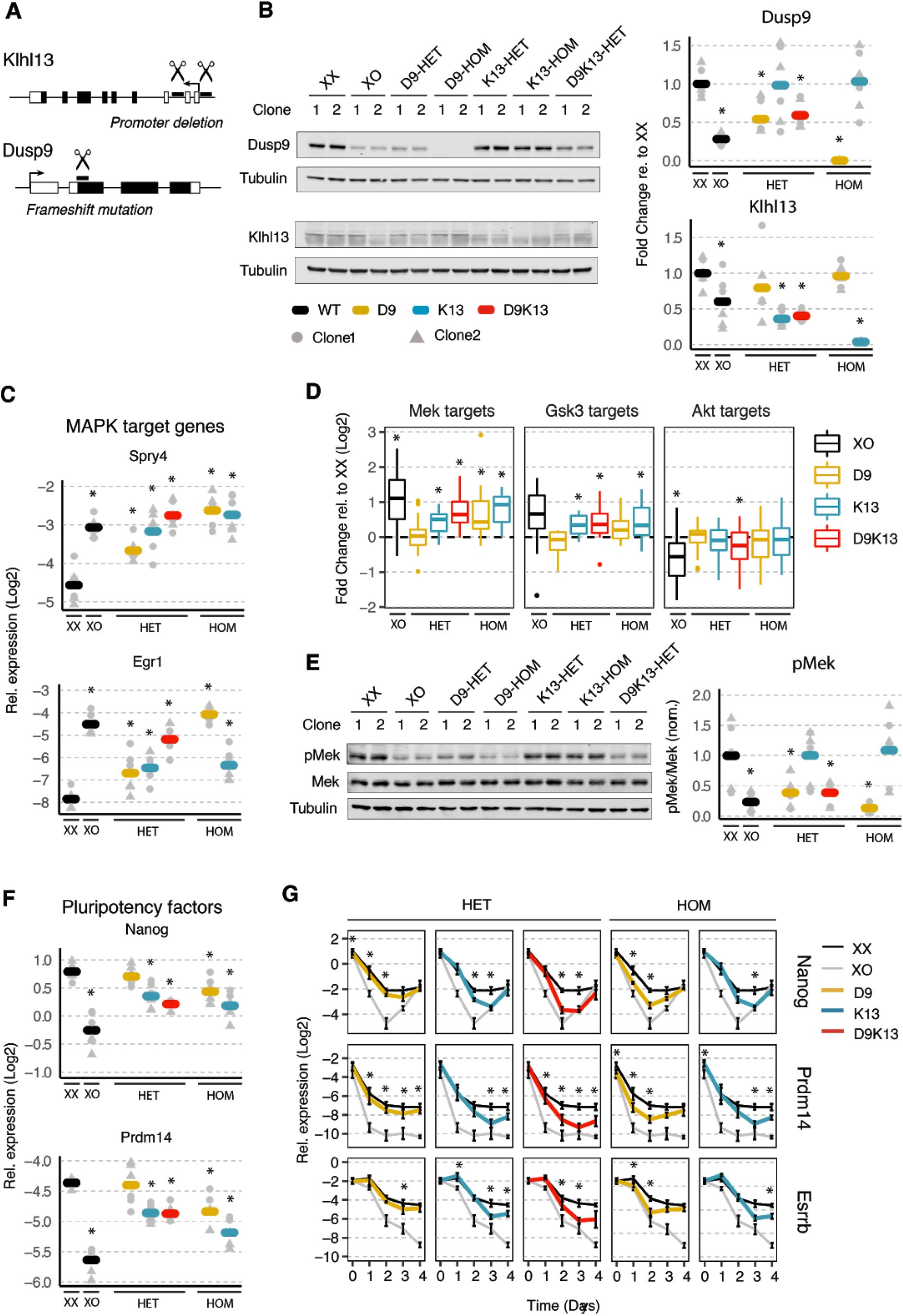
Heterozygous mutations of Klhl13 and Dusp9 in female mESCs partially phenocopy the male pluripotency state. **(A)** Schematic depiction of the strategies used to generate Klhl13 (K13) and Dusp9 (D9) mutant cell lines. **(B-F)** Comparison of female 1.8 XX mESCs with a heterozygous (HET) or homozygous (HOM) deletion of Dusp9 (yellow), Klhl13 (blue) or both (red) with the parental XX line and XO controls (2 clones per genotype). Individual measurements are shown as grey dots (clone 1) and triangles (clone 2) and the mean across two clones and three biological replicates is indicated by a thick bar. **(B)** Immunoblot quantification of Dusp9 (top) and Klhl13 (bottom) protein levels, normalized to Tubulin and to the mean of the XX controls. **(C)** Quantification of MAPK target genes by qPCR. **(D)** Boxplots showing expression of Mek (left), Gsk3 (middle) and Akt (right) target genes in cell lines with the indicated genotypes as assessed by RNA-seq. Boxes indicate the 25th to 75th percentiles and the central line represents the median. **(E)** Quantification of pMek, normalized to total Mek and to the XX control cells by immunoblotting. **(F)** Pluripotency factor expression (Nanog and Prdm14) assessed by qPCR. **(G)** qPCR quantification of pluripotency factors during differentiation by 2i/LIF withdrawal in one clone for each genotype from the cell lines used in (B-F). Mean and SD of three biological replicates is shown. * p < 0.05 Wilcoxon rank-sum test (D), otherwise two-tailed paired Student’s t-test comparing each mutant/XO cell line and XX wildtype controls.

We next investigated phosphorylation levels of Mek and observed a completely different pattern. Neither HOM nor HET mutations of Klhl13 had any effect on pMek, but levels were reduced in the Dusp9 mutants in a dose-dependent manner (Fig. 4E, Fig. S5C). The D9K13-HET double mutants resembled the D9-HET single mutants and exhibited a 2.6-fold pMek reduction compared to the wildtype XX control, thus approaching, but not reaching the 4.3-fold reduction observed in XO cells (Fig. 4E). Taken together, these results confirm the screening results that Dusp9 and Klhl13 both affect expression of MAPK target genes, but only Dusp9 has a detectable effect on Mek phosphorylation (Fig. 2G), which is in accordance with their over-expression phenotypes in male mESCs (Fig. 3). These findings are in agreement with the role of Dusp9 as an Erk phosphatase, which reduces Erk phosphorylation and consequently the Erk mediated negative feedback upstream of Mek. Although Dusp9 acts directly on the MAPK pathway, its deletion affects MAPK target gene expression less than the deletion of Klhl13 (Fig. 4D), which is in contrast to results obtained from over-expression, where Dusp9 shows stronger MAPK activation than Klhl13 (Fig. 3C).

In the next step we investigated pluripotency factor expression and differentiation kinetics. Nanog and Prdm14 expression were significantly reduced in K13-HET, but not in D9-HET lines (Fig. 4F, Fig. S5D). D9K13 double mutant cells expressed similar levels as the K13 single mutant. With a ∼1.5-fold reduction the two genes could account for ∼50% of the 2-2.4-fold decrease in Nanog and Prdm14 levels observed in XO cells (Fig. 4F). For the assessment of differentiation dynamics, cells were first adapted to 2i conditions (with Serum and LIF) for at least five passages. Since these conditions neutralize the expression differences of pluripotency factors between the cell lines in undifferentiated cells, they allow easier comparison of differentiation dynamics upon 2i/LIF withdrawal. Also here, Klhl13 had a stronger effect than Dusp9 (Fig. 4G). D9-HET mutants showed only a minimal reduction of Esrrb, Nanog and Prdm14 levels during differentiation compared to wildtype cells, while all three marker genes were reduced more in K13-HET cells (Fig. 4G). In the double D9K13 mutant the effects of the single mutants added up to nearly the levels observed in XO cells (Fig. 4G). We can conclude that Klhl13 has a stronger effect on pluripotency factor expression and differentiation than Dusp9 and that the double mutant can qualitatively, but not quantitatively reproduce the sex differences in mESC, suggesting that additional X-linked factors also contribute.

Finally we also assessed global CpG methylation with the LUMA assay (Fig. S5E). In XX control cells 31% of all CpG dinucleotides were methylated and levels were increased by ∼10% in the single HET mutants and by ∼15% in the HET double mutant and the HOM mutants. Given that D9K13-HET double mutants exhibited 44% methylation compared to 59% in XO control cells, Klhl13 and Dusp9 together could account for half of the differences seen in the XX/XO comparison.

To get a more global picture of how well the mutant lines recapitulated the XO phenotype, we performed a transcriptome comparison. For each genotype we identified autosomal genes that were differentially expressed when compared to the parental XX line. We found that 201 out of 956 differentially expressed genes (DEGs) in XO cells were also differentially expressed in K13-HET cells, but only 148 in D9-HET lines (Fig. 5A). For the D9K13 double mutant the overlap was 265 genes. A similar pattern was observed when performing principal component analysis (PCA). Also here the double mutant was found most closely to the XO controls, followed by K13-HET and D9-HET single mutants (Fig. 5B). These findings suggest that Klhl13 contributes more to X-dosage induced transcriptome changes than Dusp9, and that a combined effect of both can explain the observed sex differences best, but not completely.

**Figure 5.**
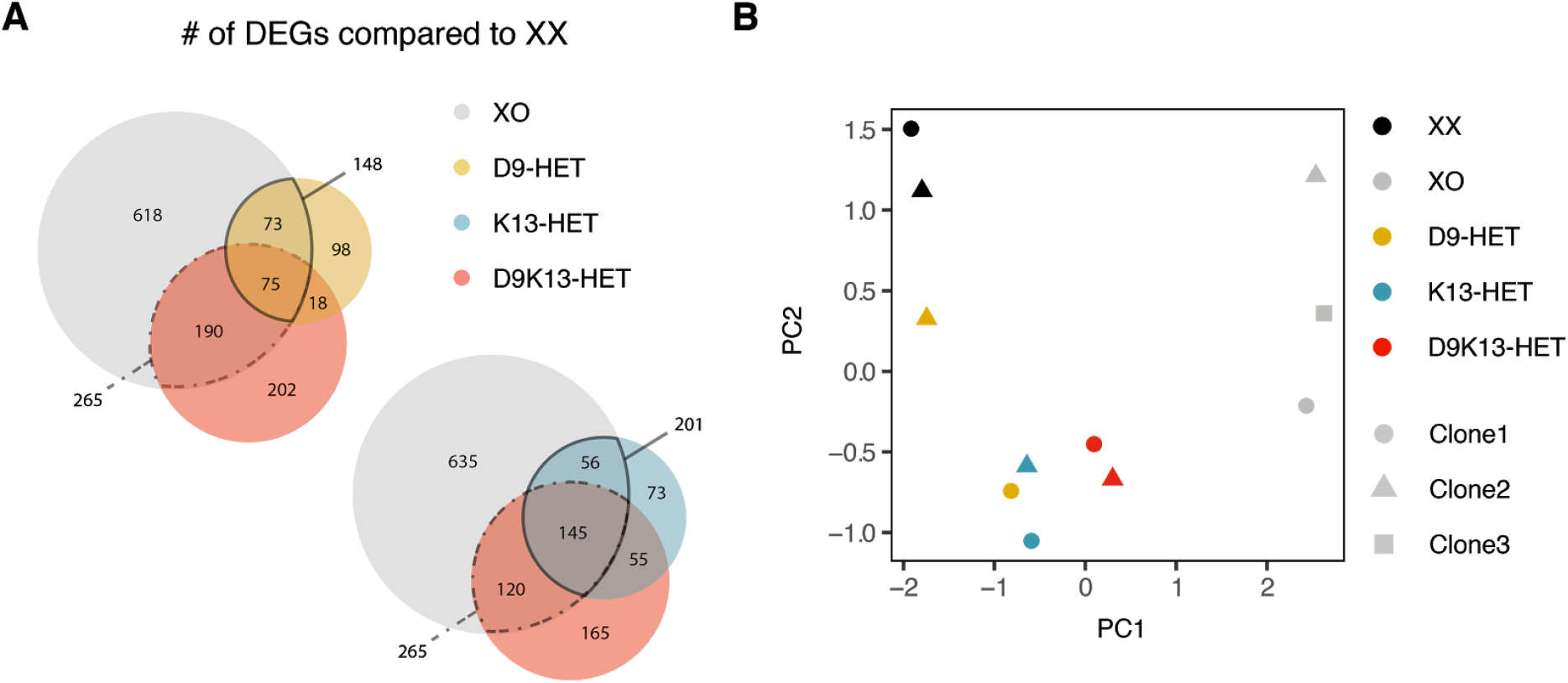
Global transcriptome profiling of Klhl13 and Dusp9 heterozygous mutant lines. **(A)** Differentially expressed genes (DEGs) in XO (grey), D9-HET (yellow), K13-HET (blue) and D9K13-HET cells (red) compared to the parental XX line were identified by RNA-seq (log_2_(fold-change) > 0.5 or log_2_(fold-change) < −0.5, p-value < 0.05). The overlap between these gene sets is shown in Venn diagrams. **(B)** Principal component analysis (PCA) of the 100 most variable genes accross XX (black), XO (grey) and heterozygous mutant cell lines (D9 yellow, K13 blue and D9K13 red), averaged across three replicates.

When comparing the results of the mutant cell lines (Fig. 4) with the over-expression experiments in male cells (Fig. 3), it becomes apparent that the relative importance of the two genes seemed to be different in the two approaches. Dusp9 had a much stronger effect than Klhl13 on MAPK target genes and pluripotency factors in the over-expression experiment, while in the mutants, both genes affected MAPK target genes, but only Klhl13 altered pluripotency factor expression. To distinguish, whether this discrepancy was due to the direction of the perturbation or different perturbations strategies used, we implemented a third validation strategy, where Dusp9 and Klhl13 were down-regulated through CRISPR interference (CRISPRi) in female mESCs, expressing an ABA inducible split KRAB-dCas9 system (Fig. 6A). For both genes, 3 different sgRNAs targeting the gene’s TSS were co-expressed from a single vector resulting in ∼20-fold reduction of mRNA expression of each gene, compared to non-targeting control sgRNAs (Fig. 6B-C). Out of 5 quantified MAPK target genes, the majority was increased upon Dusp9 and Klhl13 repression, with somewhat stronger effects for Dusp9 (Fig. 6D). The opposite pattern was observed, when profiling 5 naive pluripotency factors, where cells that downregulated Klhl13 seemed to express consistently lower levels of these genes compared to cells with Dusp9 downregulation (Fig. 6E), thus confirming the important role of Klhl13 observed with the knock-out approach (Fig. 4F).

**Figure 6.**
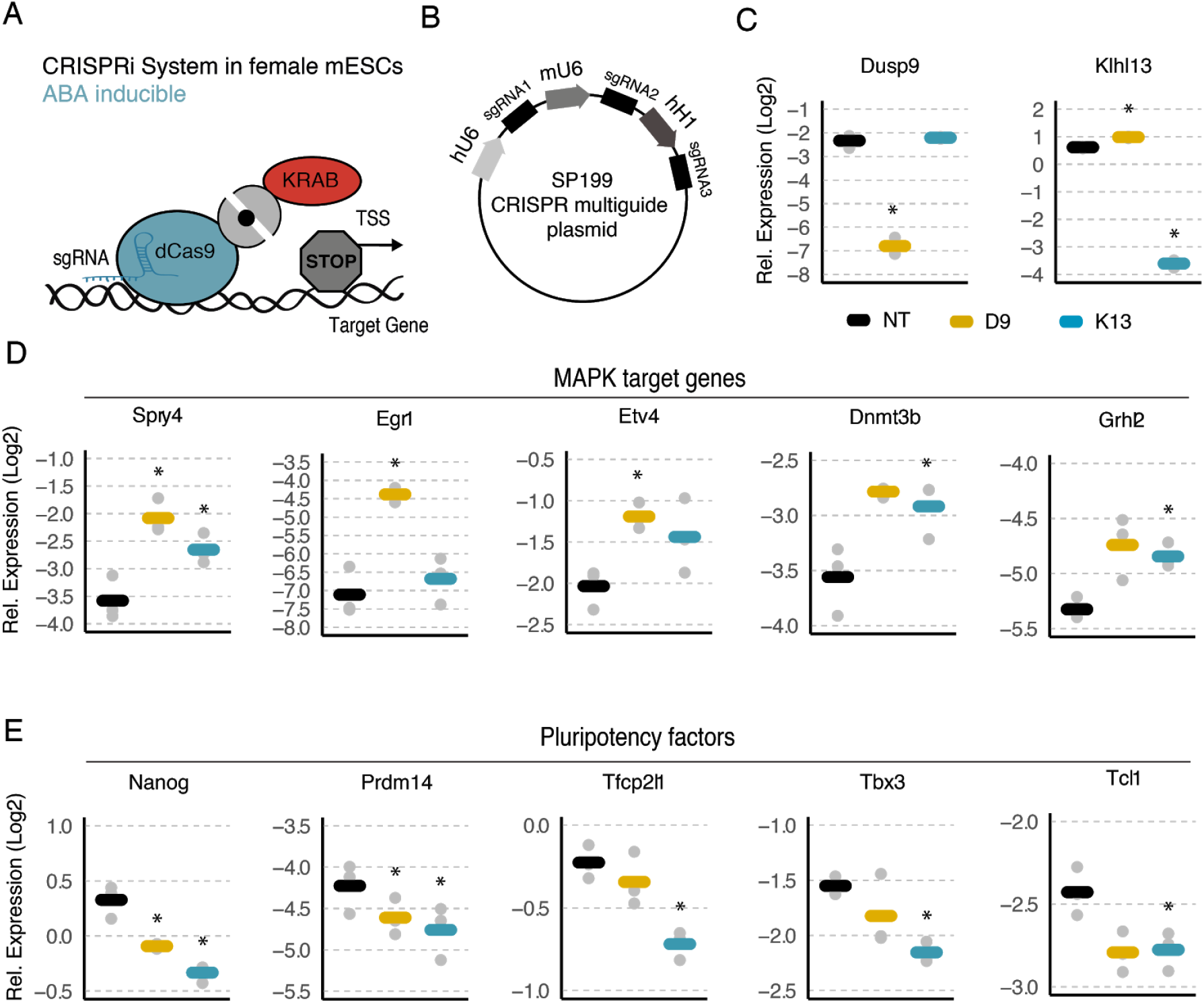
Knock-down of Dusp9 and Klhl13 in female mESCs leads to a shift towards the male pluripotency phenotype. **(A)** Catalytically dead Cas9 (blue) and the KRAB repressor domain (red) are each fused to one component of the PYL1-ABI system (grey), which dimerizes upon treatment with abscisic acid (ABA), resulting in gene repression. **(B)** CRISPR multiguide plasmid used for expression of three different sgRNAs targeting a specific gene. Each sgRNA is expressed under a different Pol III promoter, as indicated. **(C-E)** 1.8 female wildtype mESCs stably expressing the CRISPRi system shown in (A) were transduced with vectors expressing sgRNAs targeting Dusp9 (yellow), Klhl13 (blue) or a non-targeting control construct (black). Expression of each target gene (C), five MAPK target genes (D) and five pluripotency factors (E) was quantified by qPCR in cells expressing the respective sgRNAs or NTCs, as indicated. Bars represent the mean of 3 biological replicates, grey dots the individual measurements. Cells were treated with abscisic acid (ABA) for five days prior to cell harvesting for phenotypic assessment. *p < 0.05 two-tailed paired Student’s t-test comparing gene-specific sgRNAs and NTCs are indicated.

In conclusion, multiple genes underlie the female pluripotency phenotype of which we have identified and validated a novel key player, Klhl13. Dusp9 is responsible for the reduced levels of Mek phosphorylation in XX cells, but a combined effect of both genes together (partially) accounts for the global reduction of MAPK target genes in female ES cells. The pluripotency and differentiation phenotypes by contrast can primarily be attributed to reduced Klhl13 dosage in female cells. Since so far no mechanistic link between Klhl13 and pluripotency or differentiation had been reported, we set out to investigate putative Klhl13 interaction partners that might mediate the observed effects.

### Identification of Klhl13 interaction partners

Klhl13 is a member of the Cullin3 E3 ubiquitin ligase complex, where it acts as a substrate adaptor mediating protein ubiquitinylation, which might target proteins for proteasomal degradation (Fig. 7A) (Dhanoa et al. 2013; Pintard et al. 2004). We reasoned that the Klhl13-mediated sex differences we have identified might be due to reduced protein abundance of Klhl13 substrate proteins in female compared to male cells, which affect pluripotency factors, differentiation and MAPK target gene expression. To identify Klhl13 substrates in mESCs, we profiled Klhl13 interaction partners and then selected those with increased protein levels in K13-HOM mutant cells (Fig. 7B). To identify interaction partners, we ectopically expressed either full-length Klhl13 or the substrate-binding Kelch domain, tagged with a green fluorescent protein (GFP), and identified binding partners by Immunoprecipitation-Mass Spectrometry (IP-MS) using a GFP-specific antibody (Fig. 7B-D, Fig. S6A, Supp. Table S3). Since E3 ubiquitin ligases usually interact with their substrates only transiently because they are rapidly degraded, the cells were treated with a proteasomal inhibitor for their stabilization. We identified a total of 197 interaction partners for the GFP-Kelch domain and 218 for full-length Klhl13 that were enriched relative to the GFP-only controls (Fig. 7C-D, Supp. Table S3). As expected, the interaction partners identified for full-length Klhl13 and for the Kelch domain showed a large overlap, with 110 proteins being identified in both pull-downs. Two known interaction partners (Nudcd3 and Hsp90aa1) were identified with both constructs and several members of the Cullin 3 complex (Cul3, Klhl22, Klhl21, Klhl9) were found to interact with full-length Klhl13 only as expected (Sowa et al. 2009) (Fig. 7C-D, triangles).

**Figure 7.**
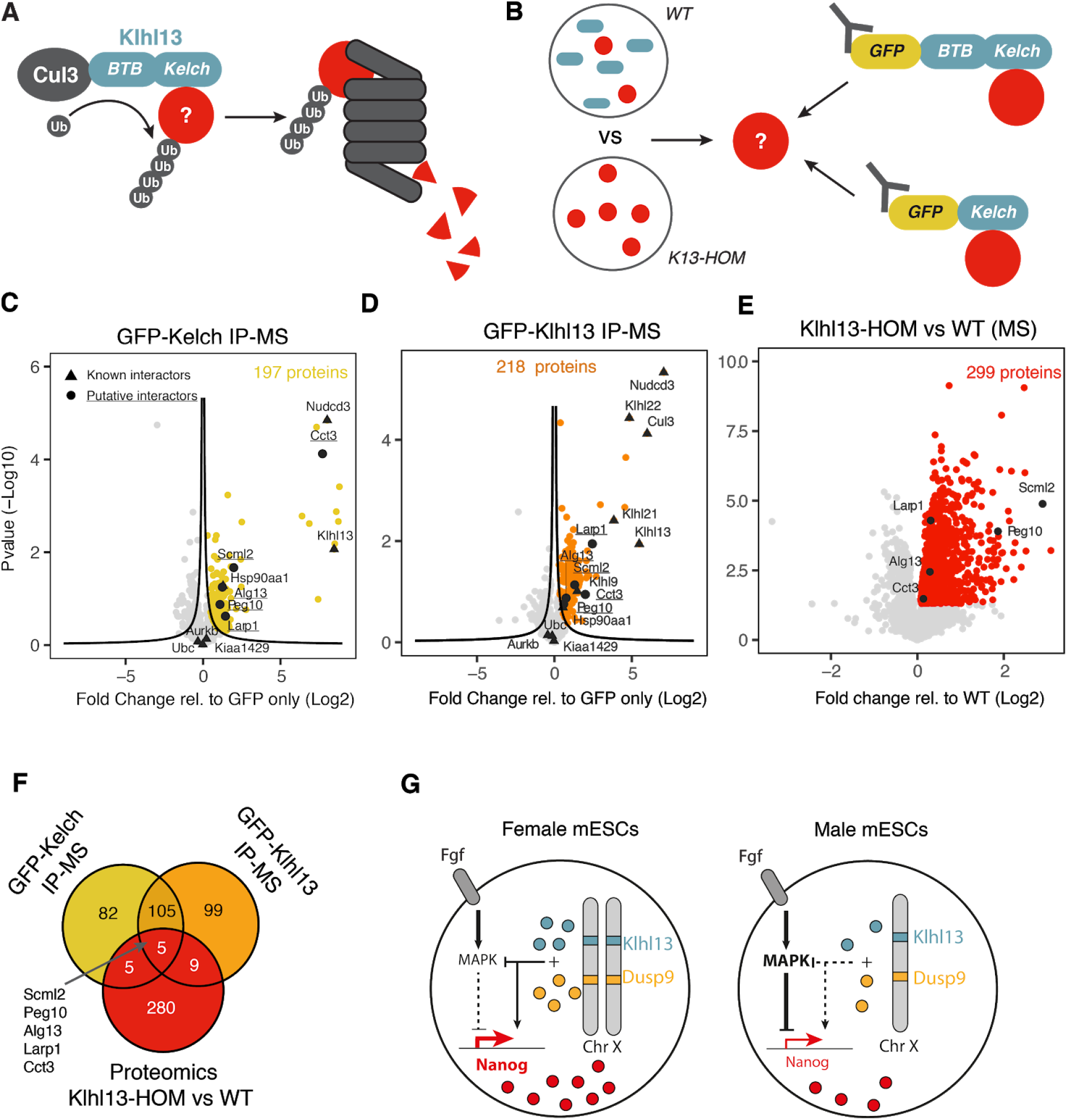
Identification of Klhl13 target proteins that mediate its effect on pluripotency and differentiation. **(A)** Schematic representation of the putative mechanism underlying Klhl13’s (blue) pluripotency-promoting effects, where differentiation-promoting substrate proteins (red) are targeted for proteasomal degradation through recruitment to the Cul3 E3 ubiquitin ligase complex via Klhl13’s Kelch domain. **(B)** Experimental strategy for the identification of Klhl13 target proteins: Putative substrates should be more abundant at the protein level in Klhl13 deficient cells and should interact with Klhl13 and with the Kelch domain only. To identify substrates the proteomes in wildtype and K13-HOM cells were compared and Klhl13/Kelch interaction partners were identified through GFP-mediated IP-MS. All three datasets were integrated to identify candidate proteins. **(C-D)** Volcano plots of the IP-MS results for the GFP-Kelch (C) and GFP-Klhl13 (D) constructs. The mean fold change across 3 biological replicates relative to the GFP-only control against the p-value calculated via a two-sample Student’s T-test with Benjamini-Hochberg correction for multiple testing is shown. Black lines indicate the significance threshold that was chosen such that FDR<0.1, assuming that all depleted proteins (left-sided outliers) were false-positive. Triangles show known Klhl13 interaction partners (BioGRID database, Supp. Table S3). **(E)** Volcano plot showing the proteome comparison of 1.8 XX wildtype cells and K13-HOM mESCs. The mean fold change across 3 biological replicates is shown. Proteins that are up-regulated upon Klhl13 depletion are highlighted in red (p<0.05 of a Student’s T-test, Benjamini-Hochberg FDR). Circles in (C-E) depict putative Klhl13 substrate proteins that were found to interact with Klhl13 and the Kelch domain and were up-regulated upon Klhl13 deletion. **(F)** Venn diagram summarizing results in C-E. **(G)** Model of how X dosage modulates the MAPK signalling pathway, pluripotency factor expression and differentiation.

To identify putative Klhl13 substrates among the 110 proteins found to interact with full-length Klhl13 and with the Kelch domain only, we quantified the total proteome of K13-HOM cells and the parental XX control line through MS with label-free quantification (Fig. 7E, Supp. Table S3). Among the 299 proteins that were significantly up-regulated in the mutant cells (p-value < 0.05, Benjamini Hochberg FDR), 5 proteins (Scml2, Peg10, Alg13, Larp1, Cct3) had been identified as putative substrates in our IP-MS experiment (Fig. 7F).

We next investigated whether any of the five identified putative Klhl13 target proteins would represent pro-differentiation factors by assessing MAPK target gene and pluripotency factor expression. To this end, we overexpressed them in female wildtype mESCs and tested whether their knock-down would rescue the phenotype of K13-HOM mutant cells. For the knock-down we used the ABA-inducible split KRAB-dCas9 (CRISPRi) described above (Fig. 6) and for gene overexpression an analogous system for gene activation, which recruits the VPR effector domains (CRISPRa) (Fig. S7A-B). Three different sgRNAs targeting the gene’s TSS were coexpressed from a single vector. Perturbation strength, as assessed by qPCR, was variable between genes, but reached at least 2-fold overexpression for all genes except Larp1 in the CRISPRa experiment and a more than 2-fold reduction for all except Cct3 upon CRISPRi (Fig. S7A-B).

We then assessed five MAPK targets (Spry4, Egr1, Etv4, Dnmt3b and Grhl2) and five naive pluripotency markders (Nanog, Prdm14, Tfcp2l1, Tbx3 and Tcl1) by qPCR (Fig. S7C-D). For factors that mediate the Klhl13 phenotype, we would expect an increase of MAPK target genes and a decrease in pluripotency markers upon over-expression, and the opposite trends upon knock-down. Generally, we only observed weak and mostly not consistent effects upon perturbation (Fig. S7C-D). However, a subset of factors exhibited some of the expected trends. Peg10 led to a small, but significant increase in MAPK target genes and downregulation of naive pluripotency factors when overexpressed in female mESCs, a trend that was confirmed in an independent experiment with another sgRNA plasmid (Fig. S7E-G). This trend was however not observed in the CRISPRi experiment. Instead, knock-down of Larp1 seemed to partially rescue the reduced pluripotency factor expression in K13-HOM cells. The reason why no effect was observed for Larp1 in the CRISPRa experiment might be its inefficient overexpression in female wildtype mESCs (Fig. S7A).

In summary, we could not identify a single gene that might mediate the effects of Khl13 on the female pluripotency phenotype through the chosen approach. Instead the phenotype might be mediated by several factors, potentially including Peg10 and Larp1. Alternatively, ubiquitinylation of Klhl13 substrates might lead to consequences other than proteasomal degradation, in which case also Klhl13 interaction partners that were not upregulated in Klhl13 knock-out cells might be involved in Klhl13 functions. We have thus narrowed down the list of candidate genes that warrant further investigation in the future.

## Discussion

We present what, to our knowledge, is the first comprehensive functional identification of genes that drive phenotypic consequences of the loss or gain of an entire chromosome. We developed a hierarchical CRISPR screening approach, which allowed us to profile a large number of genes with respect to multiple phenotypes linked to sex differences in mESCs in an unbiased manner. In an initial X-chromosome-wide screen we identified a set of candidate genes, which were then further characterized for a role in modulating three additional molecular phenotypes. In this way we identified several genes that potentially mediate X-chromosomal dosage effects and characterized the two strongest candidates in more detail, namely Dusp9 and Klhl13. Through CRISPR-mediated over-expression in male and knockout or knock-down in female cells we show that these two genes contribute to partially overlapping, yet distinct aspects of the X-dosage induced phenotype and that they appear to act in concert with additional factors. The X-dosage dependent effects in pluripotent cells can thus not be attributed to a single X-linked gene, but arise from a complex interplay of multiple factors.

Dusp9 is a phosphatase that dephosphorylates the MAPK pathway intermediate Erk and is thus a known negative regulator of the pathway (Caunt and Keyse 2013). In agreement with previous reports, we found that Dusp9 gain-of-function perturbations in male cells alter feedback strength and target gene expression (Choi et al. 2017; Li et al. 2012), while deletion of one copy of *Dusp9* in female cells results in the opposite phenotype. In addition we confirmed the previously reported alterations in global DNA methylation levels, but the magnitude of the effects was less pronounced in our study, maybe due to differences in the cell lines, culture conditions and methylation assay used (Choi et al. 2017). Dusp9 also strongly affected pluripotency factor expression and differentiation in the gain-of-function experiments, which is consistent with a previous report (Choi et al. 2017). We however observed only marginal effects in the female heterozygous Dusp9 mutant cells, again in agreement with another study (Song et al. 2019), which were however considerably stronger, when both copies of the gene were mutated or knocked-down.

For Dusp9 we thus observed a strong phenotype in the gain-of-function experiment in male cells and the opposite, albeit much weaker phenotype upon loss-of-function in female cells. Intriguingly, we found the opposite pattern for Klhl13, the second factor we investigated in detail. Here, the gain-of-function perturbation had only small effects, while loss-of-function led to an increase in MAPK target gene expression and a decrease for pluripotency factors, which was even more pronounced than the effects observed for Dusp9. This asymmetry between the gain- and loss-of-function perturbations remains puzzling and might point towards more complex interactions between multiple X-chromosomal factors.

Klhl13 is a substrate-adaptor protein of the Cul3 E3 ubiquitin ligase complex (Sumara et al. 2007) and has, to our knowledge, not yet been implicated in pluripotency, signalling or X-dosage effects. Instead, it has been reported to be involved in mitotic progression through targeting Aurora kinase B in Hela cells (Sumara et al. 2007). We could however generate mutant ES cells with a normal karyotype without difficulty, suggesting a different function for Klhl13 in ES cells. While Klhl13 did not affect phosphorylation of the MAPK pathway intermediate Mek, we found that knockout of only one copy of the gene resulted in a substantial increase in MAPK target gene expression, a reduction in pluripotency factors and more efficient differentiation.

We hypothesized that a protein, which is targeted for proteasomal degradation through Klhl13-dependent ubiquitinylation might mediate the Klhl13 phenotype. We therefore identified Klhl13-interacting proteins that were upregulated upon Klhl13 deletion. While none of the five identified proteins could fully recapitulate the Klhl13 phenotype, two of them, Peg10 and Larp1, might contribute. Peg10 is a known oncogene and has been shown to interact with Nanog and Oct4 in human cancer cells (Oliviero et al. 2015; Xie et al. 2018) and Larp1 is thought to regulate translation downstream of the mTor pathway (Fonseca et al. 2015). Identification of E3 ubiquitin ligase target proteins has also previously been reported to be challenging, probably due the transient nature of the interactions and the rapid target degradation (Iconomou and Saunders 2016). Moreover, they typically have hundreds of substrates such that the Klhl13 phenotype might be a combined effect of multiple target proteins. Another possibility is that ubiquitinylation might not lead to degradation, but might function as a signalling moiety instead (Haglund and Dikic 2005; Parvatiyar and Harhaj 2010). The Klhl13 interaction partners we have identified that were not differentially expressed in Klhl13 mutant cells, might thus warrant further investigation.

Whatever the events downstream of Klhl13 might be, or results clearly show that changes in Mek phosphorylation can be attributed completely to Dusp9, while Klhl13 appears to be the main regulator underlying the X-dosage dependent shift towards the naive pluripotent state. The combined effect of the two genes can thus account qualitatively for all aspects of the X-dosage induced phenotype. The fact that the magnitude of effects in the double-mutant does not completely reproduce those seen in XO cells, suggests a contribution of one or several additional genes. Our screening approach has identified some promising candidate genes that remain to be investigated in more detail. Moreover, additional screens in the D9K13 mutant background might allow identification of other factors in the future.

Among the other genes identified in the screens, Zic3 is a pluripotency transcription factor that induces Nanog expression and enhances iPSC generation (Lim et al. 2007, 2010; Declercq et al. 2013). Accordingly, our screen identified Zic3 as a MAPK inhibitor that promoted Nanog expression and impaired differentiation. Although it is not expressed in an X-dosage sensitive manner in the cell line we used, higher expression in female compared to male cells has been reported in other cell lines (Song et al. 2019). However, pluripotency factor expression or differentiation was found to be unaffected in heterozygous Zic3 mutant mESC lines (Song et al. 2019). Another factor identified in our screens is Stag2, a member of the Cohesin complex that has been implicated in the maintenance of pluripotency through mediating long-range regulation of pluripotency-associated genes (Kagey et al. 2010). A particularly interesting candidate is the Fthl17 gene cluster, which contains seven genes that code for ferritin-like proteins, which however lack ferroxidase activity and are partially located in the nucleus (Ruzzenenti et al. 2015). They are maternally imprinted and therefore expressed only in female, but not in male blastocysts (Kobayashi et al. 2010). Their function is completely unknown, but the fact that they exhibit a female-specific expression pattern makes them intriguing candidates that warrant a more detailed investigation. Moreover, a contribution of X-linked imprinted genes to sex differences during development is also supported by the fact that mouse embryos with an XO genotype exhibit opposite growth phenotypes compared to XX embryos depending on whether they contain the maternal or paternal X chromosome (Thornhill and Burgoyne 1993).

In summary, we report central mechanisms underlying sex differences in murine pluripotent cells. The identified genes likely contribute to the X-dosage dependent developmental delay in female embryos that has been reported in several mammalian species *in vivo* (Burgoyne et al. 1995). The X-dosage induced stabilization of the naive pluripotent state might be important to ensure that X dosage compensation has occurred before differentiation continues. Moreover, a specific differentiation speed might be required to ensure the establishment of exactly one inactive X chromosome in a female-specific manner (Mutzel et al. 2019). After having identified the relevant genes, it will now be possible to investigate the functional relevance of X-dosage effects in pluripotent cells and developmental progression, both in vitro and *in vivo*.

Since the MAPK pathway plays an important role in cancer progression, our comprehensive profiling of X-encoded MAPK regulators might lead to a better understanding of the sex bias observed in certain cancer types (Dorak and Karpuzoglu 2012). Since loss of the inactive X chromosome or partial reactivation of X-linked genes in cancer has been observed in several studies (Chaligné et al. 2015; Richardson et al. 2006), reactivation of X-linked MAPK regulators might contribute to cancer susceptibility. In the context of gender medicine, sex differences in pluripotent cells are particularly relevant for therapeutic application of iPSCs. Although conventionally cultured human iPSCs retain an inactive X chromosome, it is often eroded with passage resulting in partial, but irreversible reactivation, which will be maintained during subsequent differentiation (Vallot et al. 2015; Patel et al. 2017). Moreover, recently developed culture conditions for naive hiPS cells induce reversal of XCI (Sahakyan et al. 2017; Vallot et al. 2017). If a specific tissue is to be regenerated through *in vitro* differentiation from iPSCs, we expect that double dosage of the genes identified in our study will modulate differentiation propensity in a sex-specific manner, which has already been described for the cardiac lineage (D’Antonio-Chronowska et al. 2019). Our results will thus enable a more focussed investigation of how sex modulates iPS therapy. In conclusion, our study is a first step towards understanding how X-dosage effects shape phenotypes in a sex-specific manner.

## Material and Methods

### Molecular cloning

#### SRE-Elk reporter

For the generation of the pLenti-SRE/Elk-GFP-PEST-Hygro plasmid (Supp. Table S4), a construct consisting of the MAPK sensitive SRE-Elk promoter (containing repetitive binding sequences for the SRF (serum response factor) and Elk1 transcription factors (Sequence found in Supp. Table S4)) that drives the expression of a GFP protein fused to a destabilizing PEST sequence (kind gift from Morkel and Brummer lab) was cloned into the NheI and BsrGI (NEB) linearized hygromycin resistant lentiviral vector lenti MS2-P65-HSF1_Hygro (Addgene 61426, (Konermann et al. 2015)) by using the In-Fusion HD Cloning Kit (Takara Bio).

#### Repair template for the generation of Nanog and Esrrb reporters

Repair templates to tag Nanog and Esrrb with mCherry (pUC19-Nanog-mCherry-puro, pUC19-Esrrb-mCherry-puro, Supp. Table S4) consisted of the P2A self-cleaving peptide followed by the mCherry coding sequence and a puromycin-resistance cassette, flanked by ∼400bp homology regions to the Nanog/Esrrb locus (Esrrb-HA-Upstream: chr12:86,518,604-86,519,062, Esrrb-HA-Downstream: chr12:86,519,066-86,519,521, Nanog-HA-Upstream: chr6:122,713,142-122,713,552, Nanog-HA-Downstream: chr6:122,713,556-122,714,007 (GRCm38/mm10 Assembly)). All four fragments (upstream and downstream homology arms together with mCherry and the puromycin resistance cassette) were cloned into an XbaI (NEB) linearized pUC19 plasmid (Invitrogen) using the NEBuilder® HiFi DNA Assembly Cloning Kit (NEB) with 0.05 pmol/fragment and 10 µl of the NEBuilder master mix.

#### Klhl13 over-expression constructs

pLenti-PGK-Degron-GFP-Blast, pLenti-PGK-Degron-GFP-Klhl13-Blast, pLenti-PGK-GFP-Blast, and pLenti-PGK-GFP-Kelch, which were used to identify Klhl13 interaction partners were generated and cloned into the pLenti-PGK-GFP-Blast lentiviral plasmid (Addgene 19069, (Campeau et al. 2009)) by GenScript (Suppl. Table S4). The Klhl13 isoform expressed in mESCs was used (ENSMUST00000115313.7). The Kelch domain (AA290 to AA585) was extracted from the <u>SMART</u> (http://smart.embl-heidelberg.de/) database. The Degron sequence consists of a mutated cytosolic prolyl isomerase FKBP12^F36V^, but was not used in the reported experiments (Nabet et al. 2018). The GFP sequence was taken from previous publications (Nabet et al. 2018).

#### sgRNA design

sgRNAs targeting the Nanog, Esrrb, Klhl13, and Dusp9 locus were designed using the CRISPR-Cas9 online tool http://crispr.mit.edu:8079/. Off-target scores based on *in silico* quality and off-target predictions (Hsu et al. 2013) were compared among the candidate sgRNAs and only the top-scoring were selected.

CRISPRa sgRNA sequences targeting the TSS of the Dusp9, Cct3, Larp1, Peg10 and Scml2 genes were taken from previously published libraries (Horlbeck et al. 2016), whereas sgRNAs targeting the mESC-specific Klhl13 isoform (ENSMUST00000115313.7) were designed using the CRISPR library designer (CLD) from the Boutros lab (Heigwer et al. 2016). sgRNAs targeting the TSS of the Alg13 isoform ENSMUST00000238864.1 were designed using the CRISPOR sgRNA design tool (Concordet and Haeussler 2018).

CRISPRi sgRNA sequences targeting the TSS of the Dusp9, Alg13, Cct3, Larp1, Peg10 and Scml2 were also taken from previously published libraries (Horlbeck et al. 2016; Sanson et al. 2018), whereas sgRNAs targeting the mESC-specific Klhl13 isoform (ENSMUST00000115313.7) were designed using the CRISPOR sgRNA design tool (Concordet and Haeussler 2018). Safe-targeting control sgRNAs were implemented for multi-guide CRISPRa/CRISPRi validation experiments and sequences were extracted from previous publications (Morgens et al. 2017)

#### sgRNA cloning

For sgRNA cloning into the PX330 (PX330-Nanog_sgRNA1, PX330-Nanog_sgRNA2, PX330-Esrrb_sgRNA1, PX330-Esrrb_sgRNA2, Supp. Table S4, (Cong et al. 2013)) or PX458 plasmid (PX458-Dusp9_sgRNA1, (Ran et al. 2013)), two complementary oligos containing the guide sequence and a BbsI recognition site (Oligo F: 5’CACCGNNNNNNNNNN….3’ and Oligo R: 5’AAACNNNNNNNNNN…..C3’) were annealed and cloned into the BbsI (NEB) digested target plasmid.

sgRNAs for CRISPRa (pU6-Dusp9.1-EF1Alpha-puro-T2A-BFP, pU6-Dusp9.2-EF1Alpha-puro-T2A-BFP, pU6-Klhl13.1-EF1Alpha-puro-T2A-BFP, pU6-Klhl13.2-EF1Alpha-puro-T2A-BFP, pU6-NTC.1-EF1Alpha-puro-T2A-BFP, pU6-NTC.2-EF1Alpha-puro-T2A-BFP) were cloned into a BlpI and BstXI digested pU6-sgRNA-EF1*a*-puro-T2A-BFP plasmid (Addgene 60955, (Gilbert et al. 2014)) by annealing oligos containing the guide sequence and recognition sites for BlpI and BstXI (Oligo F: 5’TTGGNNN…NNNGTTTAAGAGC3’and Oligo R: 5’TTAGCTCTTAAACNNN…NNNCCAACAAG3’) and ligating them together with the linearized vector using the T4 DNA ligase enzyme (NEB).

For the CRISPRi validation of Dusp9 and Klhl13 (SP199_multi_Dusp9_CRISPRi and SP199_multi_Klhl13_CRISPRi, Supp. Table S4), as well as the CRISPRi/a validation of putative Klhl13 interacting partners Alg13, Cct3, Larp1, Peg10 and Scml2 (SP199_multi_NTC1, SP199_multi_NTC2, SP199_multi_Alg13_CRISPRi, SP199_multi_Cct3_CRISPRi, SP199_multi_Larp1_CRISPRi, SP199_multi_Peg10_CRISPRi, SP199_multi_Alg13_CRISPRa, SP199_multi_Cct3_CRISPRa, SP199_multi_Larp1_CRISPRa, SP199_multi_Peg10_CRISPRa, SP199_multi_Peg10_CRISPRa_2 and SP199_multi_Scml2_CRISPRa, Supp. Table S4), three different sgRNAs targeting each gene were cloned into a single sgRNA expression plasmid with Golden Gate cloning, such that each sgRNA was controlled by a different Pol III promoter and fused to the optimized sgRNA constant region described in Chen et al (Chen et al. 2013). To this end, the sgRNA constant region of the lentiGuide-puro sgRNA expression plasmid (Addgene 52963) (Sanjana et al. 2014) was exchanged for the optimized version, thus generating the vector SP199 and the vector was digested with BsmBI (NEB) overnight at 37°C and gel-purified. Two fragments were synthesized as gene blocks (IDT) containing the optimized sgRNA constant region (handle) coupled to the mU6 or hH1 promoter sequences. These fragments were then amplified with primers that contained part of the sgRNA sequence and a BsmBI restriction site (primer sequences can be found in Supp. Table S4) and purified using the gel and PCR purification kit (MachereyNagel). The vector (100 ng) and two fragments were ligated in an equimolar ratio in a Golden Gate reaction with T4 ligase and the BsmbI isoschizomer Esp3I for 20 cycles (5 min 37°C, 20 min 20°C) and transformed into NEB Stable competent E.coli (Sanjana et al. 2012). Successful assembly was verified by Sanger sequencing.

### Cell culture

#### Cell lines

Female 1.8 XX mESCs carry a homozygous insertion of 7xMS2 repeats in Xist exon 7 and are a gift from the Gribnau lab (Schulz et al. 2014). Several clones with XX or XO genotype (loss of one X chromosome) were generated through sub-cloning of the parental XX cell line. Female TX1072 ESCs, carry a doxycycline responsive promoter in front of the Xist gene on one X-chromosome and have been described previously (Schulz et al. 2014). For detailed information on the cell lines refer to Supp. Table S4. Low-passage Hek293T cells were a kind gift from the Yaspo lab.

The 1.8 SRE-Elk cell line was generated by lentiviral transduction of 1.8 XX mESCs with the pLenti-SRE/Elk-GFP-PEST-Hygro plasmid (Supp. Table S4) followed by Hygromycin (250 ng/µl, VWR) selection. Single clones were picked and expanded and GFP expression confirmed via flow cytometry.

To identify Klhl13 interactions partners, female K13-HOM mESCs (Clone 2) were transduced with the lentiviral plasmids pLenti-PGK-Degron-GFP-Blast, pLenti-PGK-Degron-GFP-Klhl13-Blast, pLenti-PGK-GFP-Blast and pLenti-PGK-GFP-Kelch plasmids (Supp. Table S4) and selected using blasticidin (5 ng/µl, Roth). Protein expression was assessed via Immunoblotting.

In 1.8-Nanog-mCherry and 1.8-Esrrb-mCherry reporter lines the C-Terminus of the coding sequences of the Nanog or Esrrb genes, respectively, is tagged with the fluorescent protein mCherry, separated by a P2A self-cleaving peptide.

Cell lines over-expressing Klhl13 and Dusp9 via the CRISPRa Suntag system were generated by lentiviral transduction of E14-STN cells, which express the CRISPR activating Sun-Tag system (Tanenbaum et al. 2014) under a doxycycline-inducible promoter (kind gift from Navarro lab, (Heurtier et al. 2019)), with plasmids carrying sgRNAs targeted to the respective promoters or non-targeting controls (pU6-Klhl13.1-EF1Alpha-puro-T2A-BFP, pU6-Klhl13.2-EF1Alpha-puro-T2A-BFP, pU6-Dusp9.1-EF1Alpha-puro-T2A-BFP, pU6-Dusp9.2-EF1Alpha-puro-T2A-BFP, pU6-NTC.1-EF1Alpha-puro-T2A-BFP, pU6-NTC.2-EF1Alpha-puro-T2A-BFP, Supp. Table S4, Fig. S3A) followed by puromycin selection (1 ng/µl, Sigma).

Dusp9 and Klhl13 heterozygous (HET) and homozygous (HOM) together with Dusp9 and Klhl13 double heterozygous mutant cell lines were generated via CRISPR/Cas9-mediated genome editing (see below) of 1.8 XX mESCs.

Cell lines for Klhl13 and Dusp9 knock-down were generated by lentiviral transduction of the 1.8 XX SP107 cell line (Clone A2, see below) with plasmids carrying sgRNAs targeting their respective promoters or a non-targeting control (SP199_multi_Dusp9_CRISPRi, SP199_multi_Klhl13_CRISPRi and SP199_multi_NTC1, Supp. Table S4). Similarly, cell lines for Alg13, Cct3, Larp1, Peg10 and Scml2 knock-down and overexpression were generated by lentiviral transduction of the 1.8 XX K13-HOM SP107 and 1.8 XX SP106 cell line (see below), respectively, with plasmids carrying sgRNAs targeting their respective promoters (SP199_multi_Alg13_CRISPRi, SP199_multi_Cct3_CRISPRi, SP199_multi_Larp1_CRISPRi, SP199_multi_Peg10_CRISPRi, SP199_multi_Alg13_CRISPRa, SP199_multi_Cct3_CRISPRa, SP199_multi_Larp1_CRISPRa, SP199_multi_Peg10_CRISPRa, SP199_multi_Peg10_CRISPRa_2 and SP199_multi_Scml2_CRISPRa, SP199_multi_NTC1 and SP199_multi_NTC2, Supp. Table S4). All cell lines were selected with puromycin (1ng/µl, Sigma) for stable sgRNA integration.

#### Cell culture and differentiation

All mESC lines were grown without feeder cells on gelatin-coated flasks (Millipore, 0.1%) in serum-containing ES cell medium (DMEM (Sigma), 15% FBS (PanBiotech), 0.1 mM ß-Mercaptoethanol (Sigma), 1000 U/ml leukemia inhibitory factor (LIF, Merck)). mESCs were passaged every second day at a density of 4×10^4^ cells/cm^2^ and medium was changed daily. Cells were differentiated by LIF withdrawal in DMEM supplemented with 10% FBS and 0.1 mM ß-Mercaptoethanol at a density of 2×10^4^ cells/cm^2^ on Fibronectin-coated dishes (Merck, 10 µg/ml).

For the differentiation of mutant cell lines (Fig. 4G), cells were first adapted to 2i+LIF medium (ES cell medium with addition of 3 μM Gsk3 inhibitor CT-99021 (Axon Medchem) and 1 μM Mek inhibitor PD0325901 (Axon Medchem)) for at least five passages before undergoing differentiation via LIF withdrawal (see above). TX1072 XX and XO cells were grown in ES cell medium supplemented with 2i and differentiated by 2i/LIF withdrawal. Hek293T cells were cultured in DMEM supplemented with 10% FBS and passaged every 2 to 3 days.

#### Lentiviral transduction

For the generation of cell lines carrying randomly integrated transgenes using lentiviral transduction, DNA constructs were first packaged into lentiviral particles. For this, 1×10^6^ Hek293T cells were seeded into one well of a 6-well plate and transfected the next day with the lentiviral packaging vectors: 1.2 µg pLP1, 0.6 µg pLP2 and 0.4 µg VSVG (Thermo Fisher Scientific), together with 2 µg of the desired construct using lipofectamine 2000 (Thermo Fisher Scientific). Hek293T supernatant containing the viral particles was harvested after 48 h. 0.2×10^6^ mESCs were seeded per 12-well and transduced the next day with 500 µl of viral supernatant and 8 ng/µl polybrene (Sigma). Antibiotic selection was started two days after transduction and kept for at least 3 passages.

#### Genome editing

To generate 1.8-Nanog-mCherry and 1.8-Esrrb-mCherry reporter lines 1×10^6^ 1.8 mESCs were transfected with 4 µg of the pUC19-Nanog-mCherry-puro or pUC19-Esrrb-mCherry-puro plasmid (Supp. Table S4) and 1.5 µg of each of the sgRNAs plasmids (PX330-Nanog-sgRNA1/2 and PX330-Esrrb-sgRNA1/2) using 16.5 µl of lipofectamine 3000 and 22 µl of P3000 (Thermo Fisher Scientific) according to manufacturer’s recommendations. Cells were selected with puromycin (1 ng/µl, Sigma) for three days, starting at day two after transfection. The puromycin selection cassette was subsequently excised by transient transfection of CRE recombinase expression plasmid pCAG-Cre (Addgene 13775, (Matsuda and Cepko 2007)). Individual clones were expanded and tested for loss of puromycin resistance. mCherry fluorescence was measured via flow cytometry and clones were subsequently genotyped by PCR (Fig. S2B). All PCRs were carried out by using the Hotstart Taq Polymerase (Qiagen), a Tm of 56°C and 30 cycles (Primer sequences are listed in Supp. Table S4).

In order to generate Klhl13 mutant mESCs, 4 guide RNAs were designed to target a 4.5 kb region around the *Klhl13* promoter (2 guide RNAs on each side) with the Alt-R® CRISPR-Cas9 System (IDT), which contains all necessary reagents for the delivery of Cas9-gRNA ribonucleoprotein complexes (RNP) into target cells. Briefly, crRNAs and tracrRNA (gRNA sequences in Supp. Table S4) were mixed in equimolar concentrations and the 4 crRNAs and tracrRNA duplexes were subsequently pooled together. 2.1 µl PBS, 1.2 µl of the tra+cr duplex (100 µM Stock), 1.7 µl Cas9 (61 µM Stock) and 1 µl electroporation enhancer were pipetted together and incubated for 20 min. 10^5^ cells were nucleofected with the mixture using the CP106 program of the Amaxa 4D-Nucleofector (Lonza) and plated on gelatin-coated 48-well plates. After 48 h, cells were seeded at a density of 10 cells/cm^2^ into 10 cm plates. Individual clones were picked, expanded and genotyped for the presence of the promoter deletion. The genotyping strategy is shown in Fig. S4B. For the amplification of the wildtype band, the HotStart Taq Polymerase (Qiagen) was used with an annealing temperature of 51°C and 35 cycles. For the deletion, the Phusion HiFi Polymerase (NEB) was used with an annealing temperature of 63°C and 35 cycles (Primer sequences are listed in Supp. Table S4).

For the generation of Dusp9 mutant mESCs, 2×10^6^ WT and K13-HET (Clone 1) cells were nucleofected with 5 µg of the PX458-Dusp9_sgRNA1 plasmid (Supp. Table S4) and subsequently plated on gelatin-coated 6cm plates. The next day, high GFP+ cells were single-cell sorted into a 96-well plate and expanded. Clones were screened for homozygous or heterozygous frameshift deletions via Sanger-sequencing and Immunoblotting. Heterozygous deletion of several selected clones was further confirmed via NGS. Briefly, a region surrounding the Dusp9 deletion was amplified using the Phusion HiFi Polymerase (NEB) with a total of 30 cycles and an annealing temperature of 65°C (Primer sequences in Supp. Table S4, OG197/OG198). A second PCR using again the Phusion HiFi Polymerase (NEB) with a total of 14 cycles and an annealing temperature of 65°C was performed in order to attach the illumina adaptors and barcodes (Supp. Table S4, OG202/OG210). A dual barcoding strategy was employed, where Illumina barcodes were included in the reverse and custom sample barcodes in the forward primers. Samples containing the same Illumina barcode but different custom sample barcodes were pooled in an equimolar fashion and sequenced on the Illumina Miseq platform PE150. Samples were aligned using Bowtie2 (Langmead and Salzberg 2012) and an index containing sample barcodes and possible deletion sequences based on previously generated sanger sequencing data, gaining approximately 4000 reads per sample.

#### Generation of cell lines expressing the KRAB/VPR-dCas9 systems using Piggybac transposition

The 1.8 XX SP107 (Clone A2) and 1.8 XX K13-HOM SP107 mESC lines stably express PYL1-KRAB-IRES-Blast and ABI-tagBFP-SpdCas9, constituting a two-component CRISPRi system, where dCas9 and the KRAB repressor domain are fused to ABI and PYL1 proteins, respectively, which dimerize upon treatment with abscisic acid (ABA) (Gao et al. 2016). The 1.8 XX SP106 mESC line, on the other hand, expressess PYL1-VPR-IRES-Blast instead of PYL1-KRAB-IRES-Blast, together with ABI-tagBFP-SpdCas9, which leads to CRISPR-mediated activation of target genes when recruited to their TSS upon ABA treatment.

The 1.8 XX SP107, 1.8 XX K13-HOM SP107 and 1.8 XX SP106 mESC lines were generated through piggybac transposition. To this end, the puromycin resistance cassettes in the piggybac CRISPRi expression plasmid pSLQ2818 (pPB: CAG-PYL1-KRAB-IRES-Puro-WPRE-SV40PA PGK-ABI-tagBFP-SpdCas9, Addgene 84241 (Gao et al. 2016)were exchanged for a blasticidin resistance, resulting in plasmid SP107 and SP106, respectively. The SP107 and SP106 plasmids were then, together with the hyperactive transposase (pBroad3_hyPBase_IRES_tagRFP) (Redolfi et al. 2019), transfected into the 1.8 XX/1.8 XX K13-HOM (Clone 1) and 1.8 XX (Clone 1) mESC lines, respectively, in a 1-to-5 transposase-to-target ratio. RFP-positive cells were sorted 24 h after transfection and cells were selected with blasticidin (5 ng/µl, Roth) for stable construct integration. After expansion, high BFP-positive cells were sorted. For the 1.8 XX SP107 mESCs a clonal line was generated. Since target-gene repression in cell lines stably expressing the SP107 construct transduced with sgRNAs was often observed already without ABA treatment, we could not make use of the inducibility of the system. Instead, 1.8 XX SP107 and 1.8 XX K13-HOM SP107 mESCs were always treated with ABA (100 µM) 5 days before the analysis and effects were compared to NTC sgRNAs. A five-day ABA treatment (100 µM) was also carried out for the 1.8 XX SP106 mESC line previous to cell harvesting.

### CRISPR KO Screens

#### sgRNA library design

sgRNA sequences were extracted from the genome-wide GeCKO library (Shalem et al. 2014). For the GeCKOx library, a list of protein-coding and miRNA genes on the X chromosome was obtained from the NCBI Reference Sequence (Refseq) track on the UCSC genome browser (Pruitt et al. 2005, 2014). Control genes were included that were annotated with the Gene Ontology (GO) terms “Erk1 and Erk2 Cascade” (GO 0070371), “Regulation of Erk1 and Erk2 Cascade” (GO 0070372), “Negative regulation of Erk1 and Erk2 Cascade” (GO 0070373) and “Positive regulation of Erk1 and Erk2 Cascade” (GO 0070374). Additionally, known MAPK regulators Grb2, Fgfr2, Dusp5, Dusp7, and Dusp2 were added as additional controls. Six sgRNAs per gene and 100 non-targeting control sgRNAs were included in the GeCKOx library. For the GeCKOxs library, the 50 most enriched and depleted X-linked genes and the 10 most enriched and depleted MAPK regulators from the primary screen were identified using HitSelect (Diaz et al. 2015). The most enriched genes were identified by comparing counts between Double-Sorted/Unsorted populations, whereas the most depleted genes were extracted by comparing counts between Unsorted/Double-Sorted populations. The 3 top-scoring sgRNAs for each gene were incorporated in the GeCKOxs library together with 10 non-targeting sgRNA controls. Additionally, 10 pluripotency regulators were added based on literature search (Sox2, Tbx3, Tcf3, Fgf2, Stat3, Esrrb, Tfcp2l1, Klf2, Nanog, Pou5f1). Klf4 was incorporated into the GeCKOxs library as a MAPK regulator (GO 0070373), having scored as a hit in the SRE/Elk screen, but was treated as a pluripotency factor control in later analyses. The sgRNA sequences are provided in Suppl. Table S1.

#### sgRNA library cloning

The GeCKOx and GeCKOxs sgRNA libraries were cloned into the lentiGuide-puro sgRNA expression plasmid (Addgene 52963, (Sanjana et al. 2014)). The vector was digested with BsmBI (NEB) overnight at 37°C and gel-purified. sgRNA sequences were synthesized by CustomArray flanked with OligoL (TGGAAAGGACGAAACACCG) and OligoR (GTTTTAGAGCTAGAAATAGCAAGTTAAAATAAGGC) sequences. For the amplification of the library, 8 or 5 (GeCKOx/GeCKOxs) PCR reactions (Primer sequences in Supp. Table S4, OG113/OG114) with approx. 5ng of the synthesized oligo pool were carried out using the Phusion Hot Start Flex DNA Polymerase (NEB), with a total of 14 cycles and an annealing temperature of 63°C in the first 3 cycles and 72°C in the subsequent 11 cycles. The amplicons were subsequently gel-purified.

Amplified sgRNAs were ligated into the vector through Gibson assembly (NEB). Two 20 µl Gibson reactions were carried out using 7 ng of the gel-purified insert and 100 ng of the vector. The reactions were pooled, EtOH-precipitated to remove excess salts which might impair bacterial transformation and resuspended in 12.5 µl H_2_O. 9 µl of the eluted DNA were transformed into 20 µl of electrocompetent cells (MegaX DH10B, Thermo Fisher Scientific) according to the manufacturer’s protocol using the ECM 399 electroporator (BTX). After a short incubation period (1h, 37°C 250rpm) in 1ml SOC medium, 9ml of LB medium with Ampicillin (0.1 mg/ml, Sigma) were added to the mixture and dilutions were plated in Agar plates (1:100, 1:1000 and 1:10000) to determine the coverage of the sgRNA libraries (600x for the GeCKOx and 2500x for the GeCKOxs). 500ml of LB media with Ampicillin were inoculated with the rest of the mixture and incubated overnight for subsequent plasmid purification using the NucleoBond Xtra Maxi Plus kit (Macherey-Nagel) following the manufacturer’s instructions. To assess library composition by deep-sequencing a PCR reaction was carried out to add illumina adaptors by using the Phusion High Fidelity DNA Polymerase (NEB), with an annealing temperature of 60°C and 14 cycles (OG125/OG126). The PCR amplicon was gel-purified by using the Nucleospin Gel and PCR Clean-up kit (Macherey-Nagel) following the manufacturer’s instructions. Libraries were sequenced paired-end 50bp on the HiSeq 2500 Platform yielding approximately 25 Mio. fragments for the GeCKOx (20 pM loading concentration) and 1.3×10^6^ fragments for the GeCKOxs library (22 pM loading concentration) (Read alignment statistics found in Supp. Table S1).

#### Viral packaging of sgRNA libraries

To generate virus carrying sgRNAs of the GeCKOx and GeCKOxs libraries, HEK293T cells were seeded into 12/8 10 cm plates and transfected the next day at 90% confluence. Each plate was transfected with 6.3 µg of pPL1, 3.1 µg of pLP2 and 2.1 µg of VSVG vectors (Thermo Fisher Scientific) together with 10.5 µg of the GeCKOx/GeCKOxs library plasmids in 1ml of Opti-MEM (Life technologies). 60 µl Lipofectamine 2000 Reagent (Thermo Fisher Scientific) were diluted in 1ml Opti-MEM. Both mixtures were incubated separately for 5 min and then combined followed by a 20 min incubation, after which they were added dropwise to the HEK293T cells. Medium was changed 6 h after transfection. Transfected HEK293T cells were cultured for 48 h at 37°C, afterwards the medium was collected and centrifuged at 1800 x g for 15 min at 4°C. Viral supernatant was further concentrated 10-fold using the lenti-X™ Concentrator (Takara Bio) following the manufacturer’s instructions and subsequently stored at −80°C.

To assess the viral titer, 5 serial 10-fold dilutions of the viral stock were applied to each well of a 6-well mESC plate (MOCK plus 10^−2^ to 10^−6^) for transduction with 8 ng/µl polybrene (Merck). Two replicates were generated for each well. Selection with puromycin (1 ng/µl, Sigma) was started two days after transduction and colonies were counted after 8 days. The number of colonies multiplied with the dilution factor yields the transducing units per ml (TU/ml), which ranged from 0.5-1.5×10^6^ TU/ml.

#### Transduction and phenotypic enrichment

For the SRE-Elk screen, female 1.8-SRE-Elk mESCs were passaged twice before transduction with viral supernatant carrying the lentiCas9 plasmid (Addgene 52962, (Sanjana et al. 2014)). Blasticidin selection (5 ng/µl, Roth) was started two days after transduction and kept for 4 passages, after which 6×10^6^ cells were transduced with the sgRNA library (MOI=0.3). Puromycin selection (1 ng/µl, Sigma) was started 48 h after transduction and kept until harvesting at day 7 after transduction. The 25% of cells with the highest reporter activity were sorted. From these cells, 6-8×10^6^ cells were snap-frozen and 6×10^6^ were cultured for two additional days and subsequently sorted for GFP (top 25%). Around 8×10^6^ unsorted cells were snap-frozen on day 7 and day 9 after transduction.

For the secondary screens, 2×10^6^ female 1.8 XX Nanog-mCherry, 1.8 XX Esrrb-mCherry (see above for description) or 1.8 XX mESCs were transduced with the lentiCas9 plasmid as described above and then with the GeCKOxs library.

1.8 XX mESCs were stained for pMek on day 7 after transduction and the 25% of cells with the lowest pMek signal were sorted. 1.8-Esrrb-mCherry mESCs were passaged for differentiation (LIF withdrawal) on day 5 and differentiated for 3 days, after which cells were harvested and the 10% cells with the lowest mCherry fluorescence were sorted. 1.8-Nanog-mCherry mESCs were harvested on day 7 and the 25% cells with the lowest mCherry fluorescence were sorted. From these cells, around 2×10^6^ were cultured for two additional days and subsequently sorted for mCherry (bottom 25%). Approximately 1×10^6^ sorted and unsorted cells were snap-frozen for subsequent library preparation from all the secondary screens in order to maintain good library representation.

#### pMek intracellular staining

For the intracellular pMek staining, colonies were washed with PBS and dissociated to single cells with a 5 min trypsin (Life technologies) incubation. Trypsinization was stopped through addition of -LIF medium. Cells were disaggregated and pelleted, washed with PBS and immediately fixed with 1.5% PFA (Roth). The cell mixture was incubated for 10 min at room temperature and subsequently centrifuged for 5 min at 500 x g.

Cells were resuspended in ice-cold MeOH, incubated for 10 min on ice (0.5 ml/1×10^6^ cells) and centrifuged for 5 min at 500 x g. Cells were washed once with staining buffer (PBS + 1% BSA (Sigma), 2 ml/1×10^6^ cells) and blocked for 10 min in staining buffer. Cells were incubated with the pMek-specific antibody (Cell Signaling, #2338,1:100, antibodies are listed in Supp. Table S4) for 30 min at room temperature (100 µl/1×10^6^ cells), then washed twice with staining buffer. Cells were then incubated with an anti-rabbit-Alexa647 antibody (Thermo Fisher Scientific,1:400) for 15 min at room temperature (100 µl/1×10^6^ cells), washed twice with staining buffer before FACS sorting using the BD FACSAria™ II.

#### Preparation of sequencing libraries

For the SRE-Elk screen, genomic DNA was isolated from the frozen cell pellets using the DNeasy Blood and Tissue kit (Qiagen) following the manufacturer’s instructions. For the secondary screens, genomic DNA from frozen cell pellets was isolated via Phenol/Chloroform extraction due to higher yields. Briefly, cell pellets were thawed and resuspended in 250 µl of Lysis buffer (1% SDS (Thermo Fisher Scientific), 0.2 M NaCl and 5 mM DTT (Roth) in TE Buffer) and incubated overnight at 65°C. 200 µg of RNAse A (Thermo Fisher Scientific) were added to the sample and incubated at 37°C for 1h. 100 µg of Proteinase K (Sigma) were subsequently added followed by a 1h incubation at 50°C. Phenol/Chloroform/Isoamyl alcohol (Roth) was added to each sample in a 1:1 ratio, the mixture was vortexed at RT for 1 min and subsequently centrifuged at 16000 x g for 10 min at room temperature. The aqueous phase was transferred to a new tube, 1 ml 100% EtOH, 90 µl 5 M NaCl and 1 µl Pellet Paint (Merck) was added to each sample, mixed, and incubated at −80°C for 1 h. DNA was pelleted by centrifugation for 16000 x g for 15 min at 4°C, pellets were washed twice with 70% EtOH, air-dried and resuspended in 50 µl H_2_O.

The PCR amplification of the sgRNA cassette was performed in two PCR steps as described previously with minor modifications (Shalem et al. 2014). In order to ensure proper library coverage (300x), each sample was amplified in 6/2 PCR reactions (2 µg DNA/reaction) in the primary/secondary screens using the ReadyMix Kapa polymerase (Roche) with a total of 20 cycles and an annealing temperature of 55°C (Primer sequences in Table S4, OG115/OG116).

Successful amplification was verified on a 1% agarose gel and a second nested PCR was performed to attach sequencing adaptors and sample barcodes with 2.5 µl of the sample from the first PCR with a total of 11 cycles and an annealing temperature of 55°C (OG125/OG126).

Resulting amplicons were loaded on a 1% agarose gel, purified using the Nucleospin Gel and PCR clean-up kit (Macherey-Nagel). Libraries from the primary screen were sequenced 2×50bp on the HiSeq 2500 Platform (18 pM loading concentration) yielding approximately 4×10^6^ fragments per sample (Read alignment statistics found in Supp. Table S1). Secondary screens were sequenced 2×75bp (Pluripotency and differentiation screens) on the Nextseq 500 (2.2 pM loading concentration) or 2×50 (pMek screen) on the HiSeq 2500 Platform (20 pM loading concentration).

#### Data Analysis

Data processing and statistical analysis was performed on the public Galaxy server usegalaxy.eu (Afgan et al. 2016) with the MAGeCK CRISPR screen analysis tools (Li et al. 2014, 2015). To this end, fastq files for read1 were uploaded to the Galaxy server. Alignment and read counting was performed with MAGeCK_count. Duplicated sgRNAs were excluded, leaving 6508 unique sgRNA sequences. Read counts and alignment statistics are given in Suppl. Table S1. Statistical analysis was performed with MAGeCK_test for each screen separately. Normalized counts and gene hit summary files were downloaded and analyzed in RStudio 3.5.3 using the stringr, tidyr, data.table, dplyr and gplots packages. For easier interpretation of the results, common names were used instead of official gene symbols for a subset of genes (Erk2, Mek1, Fthl17e, Fthl17f and H2al1m) in the figures and Supp. Table S1D. The 50 most enriched and depleted genes for the generation of the GeCKOxs sgRNA library from the primary screen were extracted using HitSelect (Diaz et al. 2015).

### DNA methylation profiling via LUMA

For the assessment of global CpG methylation levels the luminometric methylation assay (LUMA) was performed as described previously (Pilsner et al. 2010). For this, genomic DNA was isolated using the DNeasy Blood and Tissue Kit (Qiagen) and 500 ng of DNA were digested either with HpaII/EcoRI (NEB) (Tube A) or MspI/EcoRI (NEB) (Tube B) in Tango Buffer (Thermo Fisher Scientific) in a total of 20 µl for 4 h at 37°C. 15 µl of Pyrosequencing Annealing Buffer (Qiagen) were mixed with 15 µl of each sample and overhangs were quantified by Pyrosequencing using the following dispensation order GTGTGTCACACATGTGTGTG (nucleotides were pipetted in a two-fold dilution) in the PyroMark Q24 (Qiagen). The peak height from dispensation 13 (T) corresponds to the EcoRI digestion and the peak height from dispensation 14(G) corresponds to the HpaII or the MspI digestion. For each sample, the HpaII/EcoRI ratio for tube A and the MspI/EcoRI ratio for tube B were calculated. The fraction of methylated DNA is then defined as: 1 - ((HpaII/EcoRI) / (MspI/EcoRI)).

### Flow Cytometry

Cells were resuspended in Sorting buffer (1% FCS and 1 mM EDTA) or Staining Buffer (after pMek staining, PBS + 1% BSA) before flow cytometry and cells were sorted using the BD FACSAria™ II. The sideward and forward scatter areas were used for live cell gating, whereas the height and width of the sideward and forward scatters were used for doublet discrimination. Analysis of FCS files was carried out using the FlowJo V10 Software (BD Biosciences). FCS files of the gated single cell populations were visualized using RStudio and the Flowcore package.

### Immunoblotting

Lysates were prepared from ∼ 2×10^6^ cells by washing with ice-cold PBS, directly adding Bioplex Cell Lysis Buffer (Biorad) supplemented with the provided inhibitors and shaking plates at 4°C at 300 rpm for 30 min, after which the lysates were transferred to 1.5ml eppendorf tubes and centrifuged at 4°C and 4500 x g for 20 min. Protein was transferred to a clean tube and quantified using the Pierce BCA kit (Thermo Fisher Scientific).

For signaling proteins, 25 µg protein was applied per lane. For Dusp9 10 µg and for Klhl13 40 µg were loaded per lane. Proteins were transferred to Nitrocellulose membranes by using the Trans-Blot Turbo Transfer System (Biorad) under semi-dry conditions.

Membranes were blocked for 1 h with Odyssey Blocking Buffer/PBS (1:1) (Li-COR) at room temperature, followed by an incubation with primary antibody (in Odyssey Blocking Buffer/PBST (1:1)) overnight at 4°C.

Signals were detected using near-infrared dye labelled secondary antibodies and membranes were scanned using Li-COR Odyssey. Band intensities were quantified using the Image Studio Lite Ver 5.2 by calculating median intensities of the band area and subtracting the adjacent top/bottom background. Antibodies are listed in Supp. Table S4.

### RNA extraction, reverse transcription, qPCR

For gene expression profiling, ∼ 2×10^6^ cells were washed with ice-cold PBS and lysed by directly adding 500 µl of Trizol (Invitrogen). RNA was isolated using the Direct-Zol RNA Miniprep Kit (Zymo Research) following the manufacturer’s instructions. For quantitative PCR (qPCR), 1 µg RNA was reverse transcribed using Superscript III Reverse Transcriptase (Invitrogen) with random hexamer primers (Thermo Fisher Scientific) and expression levels were quantified in the QuantStudio(tm) 7 Flex Real-Time PCR machine (Thermo Fisher Scientific) using 2xSybRGreen Master Mix (Applied Biosystems) normalizing to Rrm2 and Arpo. Primer sequences are listed in Supp. Table S4.

### RNA FISH

RNA FISH was performed as described previously with minor modifications (Chaumeil et al. 2008). Briefly, cells were singled out using Accutase (Invitrogen) and placed onto Poly-L-Lysine (Sigma) coated (0.01% in H_2_O, 10 min incubation at room temperature) coverslips #1.5 (1 mm) for 10 min. Cells were fixed in 3% paraformaldehyde in PBS for 10 min at room temperature and permeabilized for 5 min on ice in PBS containing 0.5%Triton X-100 and 2 mM Vanadyl-ribonucleoside complex (New England Biolabs). Coverslips were stored in −20°C in 70% EtOH until further use.

Before incubation with the RNA probe, the fixed cells were dehydrated through an ethanol series (80, 95 and 100% twice) and subsequently air-dried. BACs purified using the NucleoBond BAC kit (Macherey-Nagel) and spanning genomic regions of HuweI (RP24-157H12) and Klhl13 (RP23-36505) were labelled by nick-translation (Abbot) using dUTP-Atto550 (Jena Bioscience) and Green dUTP (Enzo) respectively. Per coverslip, 60 ng probe was ethanol precipitated with Cot1 repeats (in order to suppress repetitive sequences in the BAC DNA that could hamper the visualization of specific signals), resuspended in formamide, denatured (10 min 75°C) and competed for 1h at 37°C. Probes were co-hybridized in hybridization buffer overnight (50% Formamide, 20% Dextran Sulfate, 2X SSC, 1 µg/µl BSA, 10 mM Vanadyl-ribonucleoside). To reduce background, three 7 min washes were carried out at 42°C in 50% Formamide/2XSSC (pH 7.2) and three subsequent 5 min washes in 2X SSC at room temperature. Cells were stained with 0.2mg/ml DAPI and mounted using Vectashield mounting medium for fluorescence (Vector Laboratories). Images were acquired using a widefield Z1 Observer (Zeiss) equipped with a 100x objective and the filter set 38 and 43 (Zeiss). Image analysis was carried out using the Zen lite 2012 software (Zeiss).

### Karyotyping

Cell lines were karyotyped via double digest genotyping-by-sequencing (ddGBS), a reduced representation genotyping method. The protocol was performed as described in the Palmers lab website, which was adapted from previously published protocols (Elshire et al. 2011). Briefly, the forward and reverse strands of a barcode adapter and common adapter were diluted and annealed, after which they were pipetted into each well of a 96-well PCR plate together with 1 µg of each sample and dried overnight (Oligo sequences are listed in Supp. Table S4). The following day samples were digested with 20 µl of a NIaIII and PstI enzyme mix (NEB) in NEB Cutsmart Buffer at 37°C for 2 h.

After the digest, a 30 µl mix with 1.6 µl of T4 DNA ligase (NEB) was added to each well and placed on a thermocycler (16°C 60 min followed by 80°C 30 min for enzyme inactivation). By doing this, barcode and common adapters with ends complementary to those generated by the two restriction enzymes were ligated to the genomic DNA.

Samples were cleaned with CleanNGS beads (CleanNA) using 90 µl of beads for each well and following manufacturers instructions. Samples were eluted in 25 µl ddH_2_O and DNA was quantified using a dsDNA HS Qubit assay (Thermofisher). Samples were pooled in an equimolar fashion, size-selected (300-450bp) by loading 400 ng of each pooled sample on an agarose gel followed by a

cleaning step using the Nucleospin Gel and PCR Cleanup kit (Macherey-Nagel). Samples were PCR amplified using the Phusion High-Fidelity DNA Polymerase (NEB) and an annealing temperature of 68°C over 15 amplification cycles (OG218/OG219). Resulting amplicons were cleaned with CleanNGS beads in a 1:1.2 ratio (sample:beads) and sequenced with 2×75bp on the Miseq platform (12 pM loading concentration), yielding from 0.2×10^6^ to 1×10^6^ fragments per sample.

Data processing and statistical analysis was performed on the public Galaxy server usegalaxy.eu. Briefly, fastq files were uploaded and demultiplex using the “Je-demultiplex” tool (Girardot et al. 2016). Reads were mapped to the mm10 mouse reference genome (GRCm38) using “Map with BWA” (Li and Durbin 2009, 2010). Read counts for each chromosome were calculated with “multiBamSummary” (Ramírez et al. 2016) and normalized to a previously karyotyped XX control cell line (using dapi stained metaphase spreads and chromosome painting).

### RNA-seq

For the RNA-sequencing of 1.8 XX and 1.8 XO cell lines (Fig. 2), libraries were generated using the Tru-Seq Stranded Total RNA library preparation kit (Illumina) with 1 µg starting material and amplified with 15 Cycles of PCR. Libraries were sequenced 2×50bp on one HiSeq 2500 lane (22 pM loading concentration), which generated ∼40 Mio. fragments per sample. The reads were mapped with the STAR aligner allowing for maximally 2 mismatches to the mm10 mouse reference genome (GRCm38) and quantified using the ENSEMBL gene annotation (Dobin et al. 2013). Gene expression values (rpkm) were obtained using the EdgeR package in RStudio (Robinson et al. 2010).

For RNA-sequencing of the mutant cell lines (Fig. 4) the QuantSeq 3’ mRNA-Seq Library Prep Kit (FWD) for Illumina (Lexogen) was used with 800 ng starting material. Samples were sequenced with 1×75bp on the NextSeq 500 Platform (2 pM loading concentration). The count matrix was generated with the FWD-UMI Mouse (GRCm38) Lexogen QuantSeq 2.6.1 pipeline from the BlueBee NGS data analysis platform (https://www.bluebee.com/). Differential expression analysis was carried out using the EdgeR package in RStudio, together with normalization of gene expression values (cpm) (Robinson et al. 2010). Alignment statistics and normalized RNA-seq data from both datasets is provided in Supp. Table S2.

### Single-Cell RNA-seq data analysis

For reanalysis of previously published scRNA-seq data the normalized counts and the cell type annotation were downloaded from https://github.com/rargelaguet/scnmt_gastrulation. Sex annotation was provided by Ricard Argelaguet. For comparison of individual genes between male and female cells, a Wilcox ranksum test was performed using the wilcox.test function in R. For comparing chromosome-wide expression, counts for all genes located on a specific chromosome were summed up for each cell and then compared with a Wilcox ranksum test as described above. For the analysis of gene groups (naive and primed pluripotency markers) the log2-transformed counts for all genes in the group were averaged for each cell and then analyzed as above.

### Mass Spectrometry

#### GFP Immunoprecipitation

The GFP Immunoprecipitation protocol was performed as described previously with minor modifications (Hubner et al. 2010). Briefly, cells were treated with 15 µM of MG132 for 3 h prior to harvesting. Cells were pelleted and resuspended in 1 ml of Lysis Buffer containing 150 mM NaCl, 50 mM Tris, pH 7.5, 5% glycerol, 1% IGEPAL-CA-630, 1 mM MgCl_2_, 200 U benzonase (Merck), and EDTA-free complete protease inhibitor cocktail (Roche). Cells were incubated on ice for 30 min to

allow cell lysis. Lysates were centrifuged at 4,000 x g and 4°C for 15 min and the supernatant was incubated with 50 µl magnetic beads coupled to monoclonal mouse anti-GFP antibody (Miltenyi Biotec) for 20 min on ice. Magnetic columns were equilibrated by washing first with 250 µl of 100% EtOH followed by two washes with the same volume of lysis buffer. After the 20 min incubation, the lysates were applied to the column followed by three washes with 800 µl of ice-cold wash buffer I (150 mM NaCl, 50 mM Tris, pH 7.5, 5% glycerol, and 0.05% IGEPAL-CA-630) and two washes with 500 µl of wash buffer II (150 mM NaCl, 50 mM Tris, pH 7.5, and 5% glycerol). Column-bound proteins were subsequently pre-digested with 25 µl 2 M urea in 50 mM Tris, pH 7.5, 1 mM DTT, and 150 ng trypsin (Roche) for 30 min at room temperature. Proteins were eluted by adding two times 50 µl elution buffer (2 M urea in 50 mM Tris, pH 7.5, and 5 mM chloroacetamide). Proteins were further digested overnight at room temperature. The tryptic digest was stopped by adding formic acid to a final concentration of 2%.

#### Sample Preparation for proteomics with Label-Free Quantification (LFQ)

Proteomics sample preparation was done according to a published protocol with minor modifications (Kulak et al. 2014). Approximately 2×10^7^ cells were lysed under denaturing conditions in a buffer containing 3 M guanidinium chloride (GdmCl), 5 mM tris(2-carboxyethyl)phosphine, 20 mM chloroacetamide and 50 mM Tris-HCl pH 8.5. Lysates were denatured at 95°C for 10 min shaking at 1000 rpm in a thermal shaker and sonicated in a water bath for 10 min. A small aliquot of cell lysate was used for the bicinchoninic acid (BCA) assay to quantify the protein concentration. 50 µg protein of each lysate was diluted with a dilution buffer containing 10% acetonitrile and 25 mM Tris-HCl, pH 8.0, to reach a 1 M GdmCl concentration. Then, proteins were digested with LysC (Roche, Basel, Switzerland; enzyme to protein ratio 1:50, MS-grade) shaking at 700 rpm at 37°C for 2h. The digestion mixture was diluted again with the same dilution buffer to reach 0.5 M GdmCl, followed by a tryptic digestion (Roche, enzyme to protein ratio 1:50, MS-grade) and incubation at 37°C overnight in a thermal shaker at 700 rpm.

#### LC-MS/MS Instrument Settings for Shotgun Proteome Profiling

Peptide desalting was performed according to the manufacturer’s instructions (Pierce C18 Tips, Thermo Scientific, Waltham, MA). Desalted peptides were reconstituted in 0.1% formic acid in water and further separated into four fractions by strong cation exchange chromatography (SCX, 3M Purification, Meriden, CT). Eluates were first dried in a SpeedVac, then dissolved in 5% acetonitrile and 2% formic acid in water, briefly vortexed, and sonicated in a water bath for 30 seconds prior injection to nano-LC-MS. LC-MS/MS was carried out by nanoflow reverse phase liquid chromatography (Dionex Ultimate 3000, Thermo Scientific) coupled online to a Q-Exactive HF Orbitrap mass spectrometer (Thermo Scientific), as reported previously (Gielisch and Meierhofer 2015). Briefly, the LC separation was performed using a PicoFrit analytical column (75 μm ID × 50 cm long, 15 µm Tip ID; New Objectives, Woburn, MA) in-house packed with 3-µm C18 resin (Reprosil-AQ Pur, Dr. Maisch, Ammerbuch, Germany). Peptides were eluted using a gradient from 3.8 to 38% solvent B in solvent A over 120 min at 266 nL per minute flow rate. Solvent A was 0.1 % formic acid and solvent B was 79.9% acetonitrile, 20% H_2_O, 0.1% formic acid. For the IP samples, a one hour gradient was used. Nanoelectrospray was generated by applying 3.5 kV. A cycle of one full Fourier transformation scan mass spectrum (300-1750 m/z, resolution of 60,000 at m/z 200, automatic gain control (AGC) target 1×10^6^) was followed by 12 data-dependent MS/MS scans (resolution of 30,000, AGC target 5×10^5^) with a normalized collision energy of 25 eV. In order to avoid repeated sequencing of the same peptides, a dynamic exclusion window of 30 sec was used. In addition, only peptide charge states between two to eight were sequenced.

#### Data Analysis

Raw MS data were processed with MaxQuant software (v1.6.0.1) and searched against the mouse proteome database UniProtKB with 22,286 entries, released in December 2018. Parameters of MaxQuant database searching were a false discovery rate (FDR) of 0.01 for proteins and peptides, a minimum peptide length of seven amino acids, a first search mass tolerance for peptides of 20 ppm and a main search tolerance of 4.5 ppm, and using the function “match between runs”. A maximum of two missed cleavages was allowed for the tryptic digest. Cysteine carbamidomethylation was set as fixed modification, while N-terminal acetylation and methionine oxidation were set as variable modifications. Contaminants, as well as proteins identified by site modification and proteins derived from the reversed part of the decoy database, were strictly excluded from further analysis.

The calculation of significantly differentially expressed proteins for both the proteomics (K13 HOM vs XX wildtype) as well as the IP datasets (GFP-Kelch/D-GFP-Klhl13 vs GFP/D-GFP) was done with Perseus (v1.6.1.3). LFQ intensities, originating from at least two different peptides per protein group were transformed by log_2_. Only groups with valid values in at least one group were used, missing values were replaced by values from the normal distribution. Statistical analysis for differential expression was done by a two-sample t-test with Benjamini-Hochberg (BH, FDR of 0.05) correction for multiple testing. The processed output files can be found in Supp. Table S3.

For the identification of Klhl13 interaction partners, cut-offs were set from the data displayed in the volcano plots using a previously published method (Keilhauer et al. 2015). Briefly, a graphical formula as a smooth combination of the following parameters was implemented:

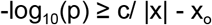

x: enrichment factor of a protein

p: p-value of the t-test, calculated from replicates

x_o_: fixed minimum enrichment

c: curvature parameter

We optimized parameters c and xo such as to have 10% FDR (left-sided outliers) while maximizing the number of right-sided outliers. In the case of the GFP-Kelch IP, c = 0.32 and x_o_ = 0.02. For the GFP-Klhl13 IP, c = 0.28 and x_o_ = 0.04. Proteins without an associated gene name were filtered out in further analyses.

## Data availability

The data sets generated during this study are available via GEO with identifiers GSE142348, GSE142349, GSE142350 (SuperSeries GSE143784) and via ProteomeXchange with identifiers PXD016729 and PXD017875. Reviewers can access the data as follows: https://www.ncbi.nlm.nih.gov/geo/query/acc.cgi?acc=GSE142348 (Token: eluxgqiwfxolnav) https://www.ncbi.nlm.nih.gov/geo/query/acc.cgi?acc=GSE142349 (Token: ehudsuekzxkrviz) https://www.ncbi.nlm.nih.gov/geo/query/acc.cgi?acc=GSE142350 (Token: uvktuecidbcthch) https://www.ebi.ac.uk/pride/ (Username/, Password: AKiKQG9S)

## Acknowledgements

We would like to thank Nils Blüthgen, Heiner Schrewe and Petra Knaus for the fruitful discussions; Joost Gribnau for providing the 1.8 XX cell line; Pablo Navarro for providing the E14 SunTag cell line; Jörn Schmiedel for sharing the sgRNA and repair plasmids for the generation of the Nanog and Esrrb reporter cell lines; Vincent Pasque for sharing Scml2 over-expressing plasmids; Stephen Keyse for providing the anti-Dusp9 antibody; Luca Giorgetti for sharing the Piggybac plasmid; Ricard Argelaguet for sharing the sex annotation of the scRNA-seq data set and the sequencing, mass spectrometry and flow cytometry facility teams at the Max Planck Institute for molecular genetics, specially Dr. Bernd Timmermann, Dr. David Meierhofer, Sven Klages, Norbert Mages, Daniela Roth, Ilona Witzke, Myriam Hochradel, Beata Lukaszewsa-McGreal, Dr. Claudia Giesecke-Thiel and Uta Marchfelder. This work was supported by the Max-Planck Research Group Leader program, E:bio Module III—Xnet grant (BMBF 031L0072) and Human Frontiers Science Program (CDA-00064/2018) to E.G.S. O.G. is supported by the DFG (GRK1772, Computational Systems Biology).

## Author Contributions

EGS and OG conceived the present work and planned the experiments. OG, AAM and ID carried out the experiments. MB contributed to experimental planning and interpretation of results concerning the CRISPR knockout screens. EGS and OG took the lead in writing the manuscript. All authors provided critical feedback and helped shape the research, analysis and manuscript.

## Supplemental Material

Supplemental Figure S1

Supplemental Figure S2

Supplemental Figure S3

Supplemental Figure S4

Supplemental Figure S5

Supplemental Figure S6

Supplemental Figure S7

Supplemental Table S1

Supplemental Table S2

Supplemental Table S3

Supplemental Table S4

---

**Supplementary Figure 1.**
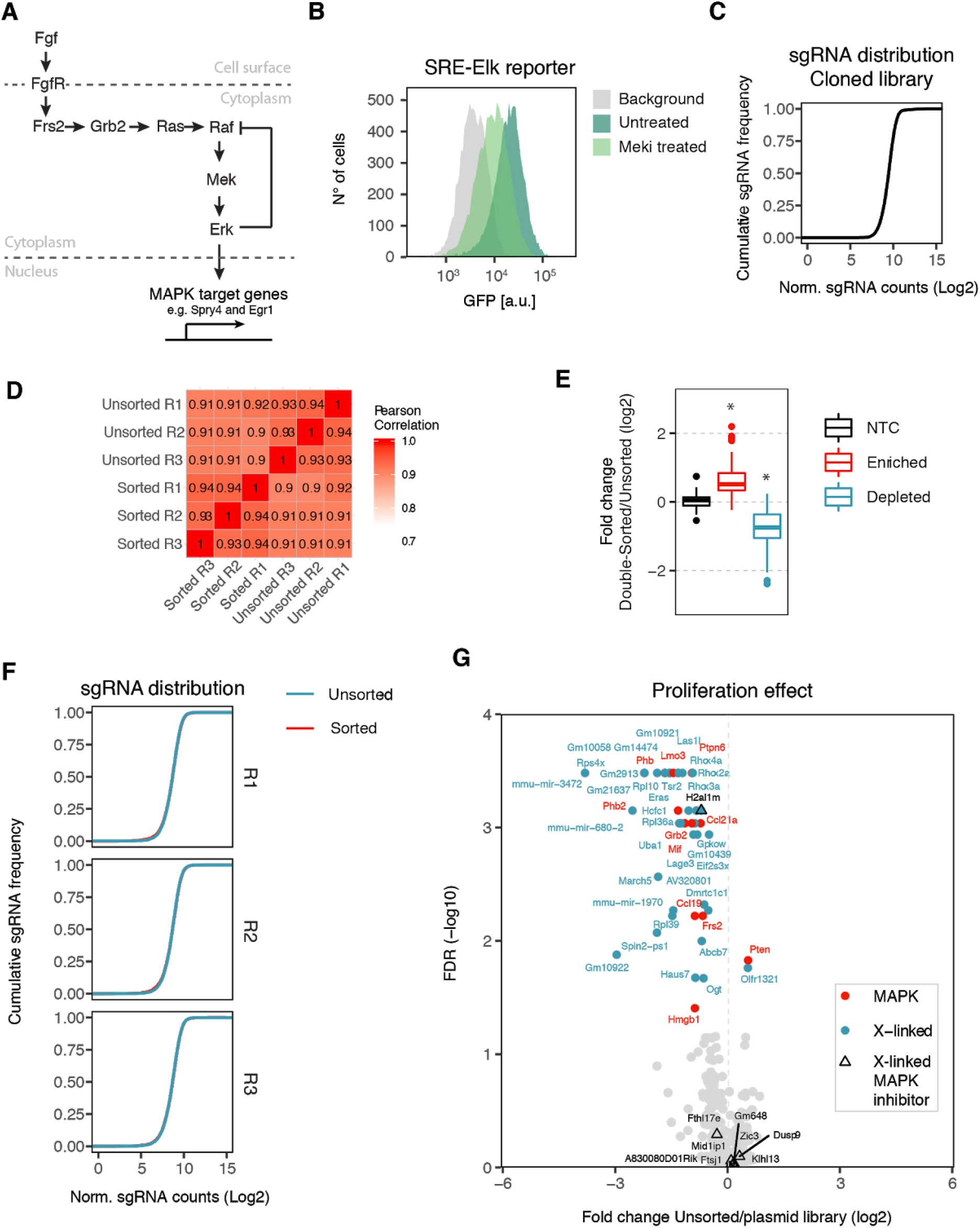
Identification of X-chromosomal MAPK regulators through a pooled CRISPR knockout screen. **(A)** Schematic description of the MAPK signaling pathway. Briefly, the fibroblast growth factor (Fgf) binds and activates the Fgf receptor (FgfR), leading to the formation of a complex consisting of the FgfR Substrate 2 (Frs2) and growth factor receptor-bound protein 2 (Grb2). This complex activates the small GTPase Ras which triggers the kinase cascade of Raf, Mek and Erk. Phosphorylated Erk translocates to the nucleus and activates MAPK target genes. Erk-dependent deactivation of Raf constitutes a strong negative feedback loop. **(B)** Flow cytometry measurement of GFP fluorescence in 1.8-SRE-Elk ESCs, treated for 48 h with 1 µM of the Mek inhibitor U0126 (Meki treated) or with DMSO (Untreated). The parental 1.8 mESC line is shown in grey. **(C)** sgRNA distribution in the cloned GeCKOx library. **(D)** Heatmap showing the pearson correlation coefficients between the sgRNA counts of all unsorted and double-sorted fractions in the MAPK screen. **(E)** Mean fold change between the double-sorted and unsorted populations for non-targeting controls and for individual sgRNAs targeting significantly enriched or depleted genes (FDR < 0.05, MAGeCK), * p < 0.05, Wilcoxon rank-sum test. **(F)** sgRNA distribution in sorted and unsorted fractions in all 3 replicates. **(G)** To detect genes that affect growth of mESCs, the sgRNA abundance was compared in the unsorted cells (day 7 after transduction) and the cloned plasmid GeCKOx library. Genes with a FDR<0.05 (MAGeCK) are highlighted in red (MAPK controls) and blue (X-linked genes). Triangles mark the identified X-linked MAPK inhibitors (compare Fig. 1C).

**Supplementary Figure 2.**
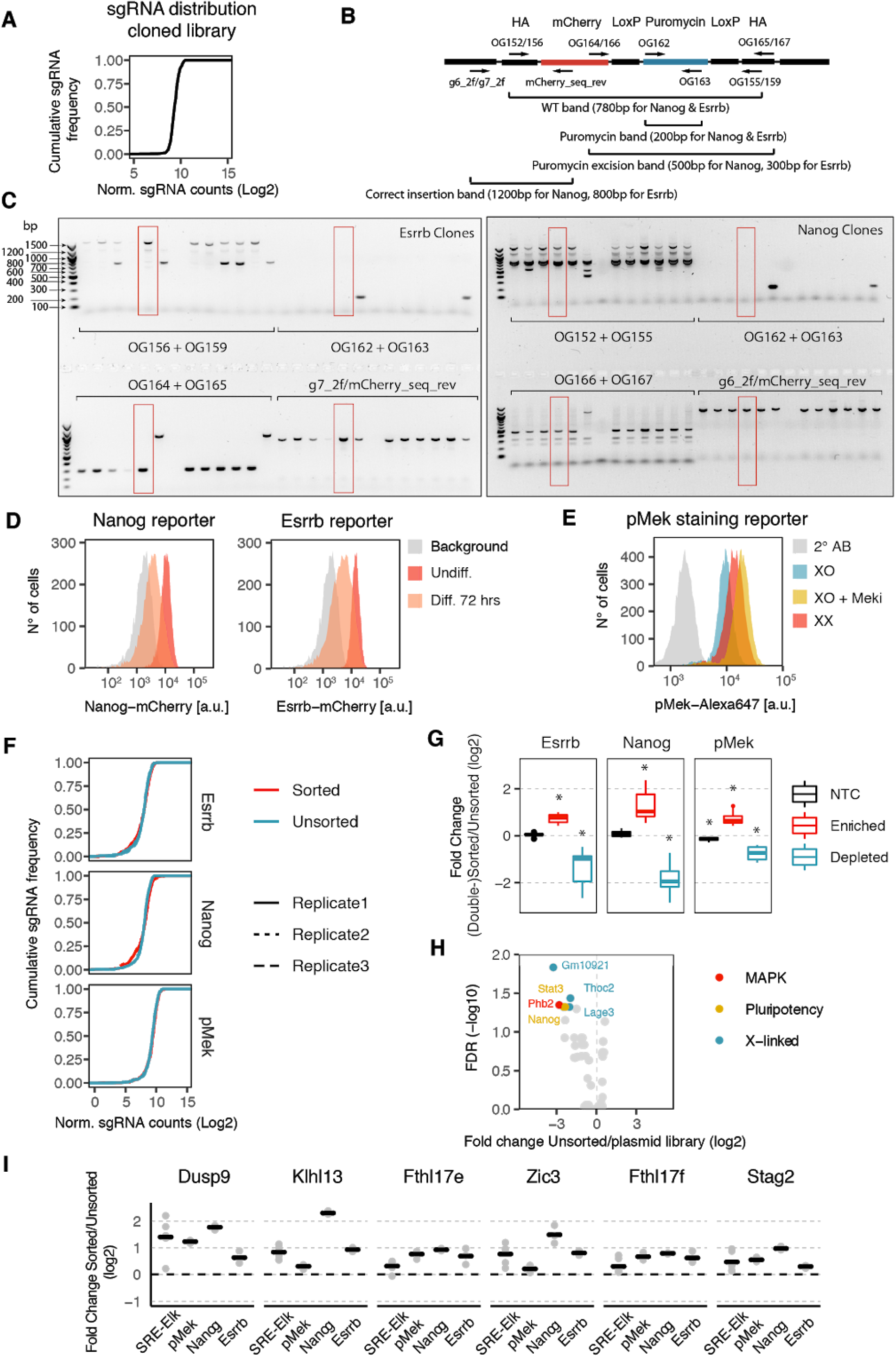
Secondary CRISPR screens profiling pluripotency factors, differentiation kinetics and Mek phosphorylation. **(A)** sgRNA distribution of the cloned GeCKOxs library. **(B-C)** Genotyping of Nanog and Esrrb tagged cell lines. **(B)** Primer locations are shown as arrows and are labelled with the primer number. The expected amplicons are shown below. **(C)** Genotyping PCR results using the indicated primer pairs, clones used for the secondary screens are highlighted in red. **(D)** Flow cytometry measurement of 1.8-Nanog-mCherry (left) and 1.8-Esrrb-mCherry mESCs (right) before and after 3 days of differentiation, showing the expected differentation-induced down-regulation of the two pluripotency factors. **(E)** Intracellular pMek staining of 1.8 XX, XO and XO cells treated with Mek inhibitor (Meki) PD0325901 for 48 h, showing the expected feedback-mediated increase in XX mESCs and upon Meki treatment. **(F)** Cumulative sgRNA distribution in unsorted and sorted fractions for all replicates of all 3 secondary screens. **(G)** Mean fold change between the (double-)sorted and unsorted populations for non-targeting controls and individual sgRNAs targeting significantly enriched or depleted genes (* p < 0.05, Wilcoxon rank-sum test). **(H)** To detect genes that affect growth of mESCs, the sgRNA abundance was compared in the unsorted cells (day 7 after transduction) and the cloned plasmid GeCKOxs library. Genes with a FDR<0.05 (MAGeCK) are highlighted. **(I)** Screen results for all individual sgRNAs targeting enriched candidate genes shown in main Fig. 2H.

**Supplementary Figure 3.**
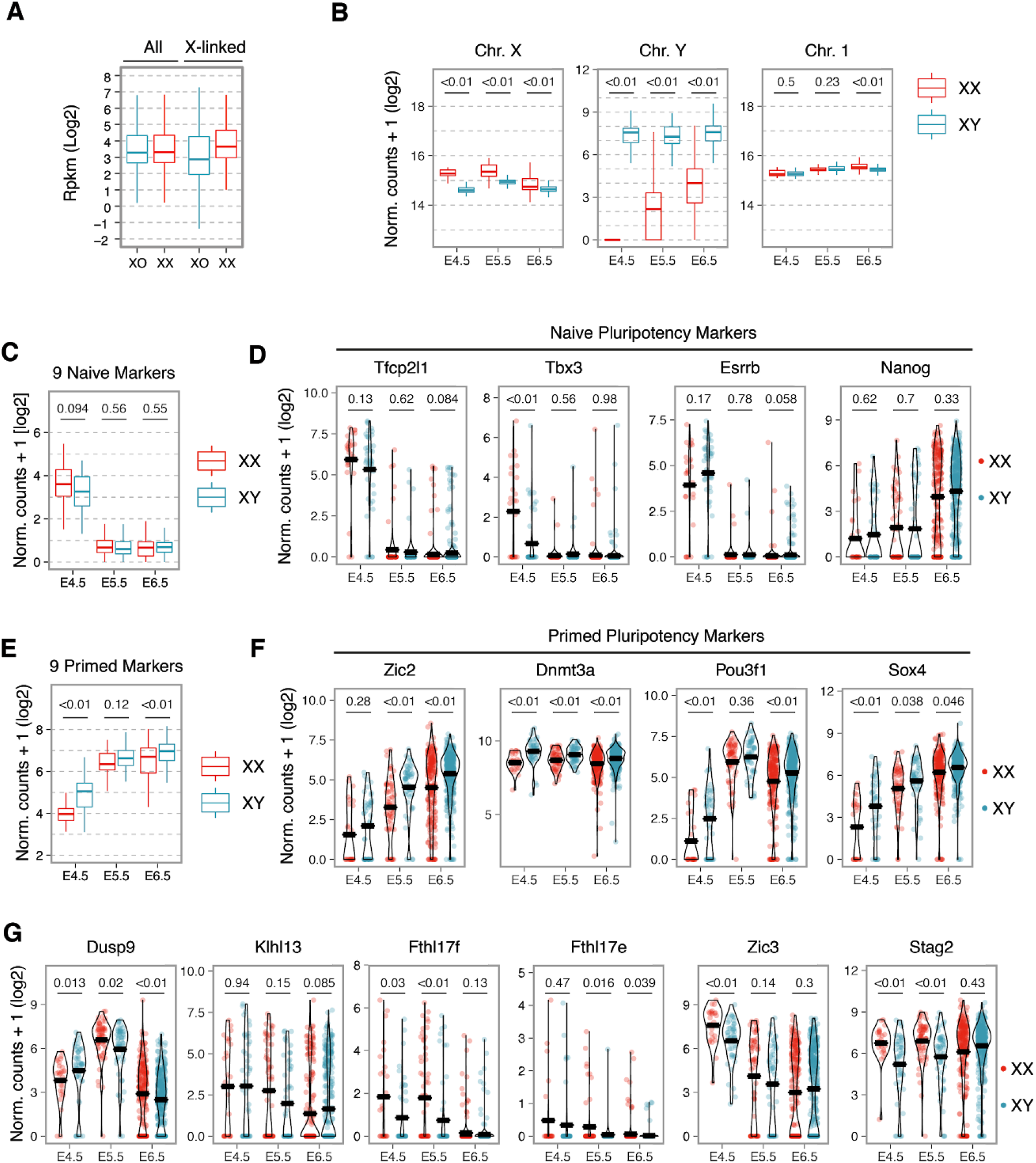
Sex differences in embryonic stem cells and mouse embryos. **(A)** Boxplot showing expression levels of all genes and only X-linked genes in 1.8 XX and 1.8 XO mESCs. **(B-G)** Comparison of male and female epiblast cells at three time points of embryonic development by single-cell RNA sequencing (Argelaguet et al). Expression in individual cells are shown for entire chromosomes (B), mean expression for each cell of 9 naive pluripotency factors (Tfcp2l1, Tbx3, Esrrb, Rex1, Klf2, Klf5, Tcl1, Nanog, Klf4) (C), expression of 4 naive pluripotency factors (D), mean expression for each cell of 9 primed pluripotency factors (Fgf5, Otx2, Pou3f1, Dnmt3b, Lef1, Sox4, Zic2, Sall2, Dnmt3a) (E), expression of 4 primed pluripotency factors (F) are shown and expression of X-linked genes identified through CRISPR screening (G). Thick bars in D,F,G indicate the mean. p-values for a Wilcoxon rank sum test are indicated.

**Supplementary Figure 4.**
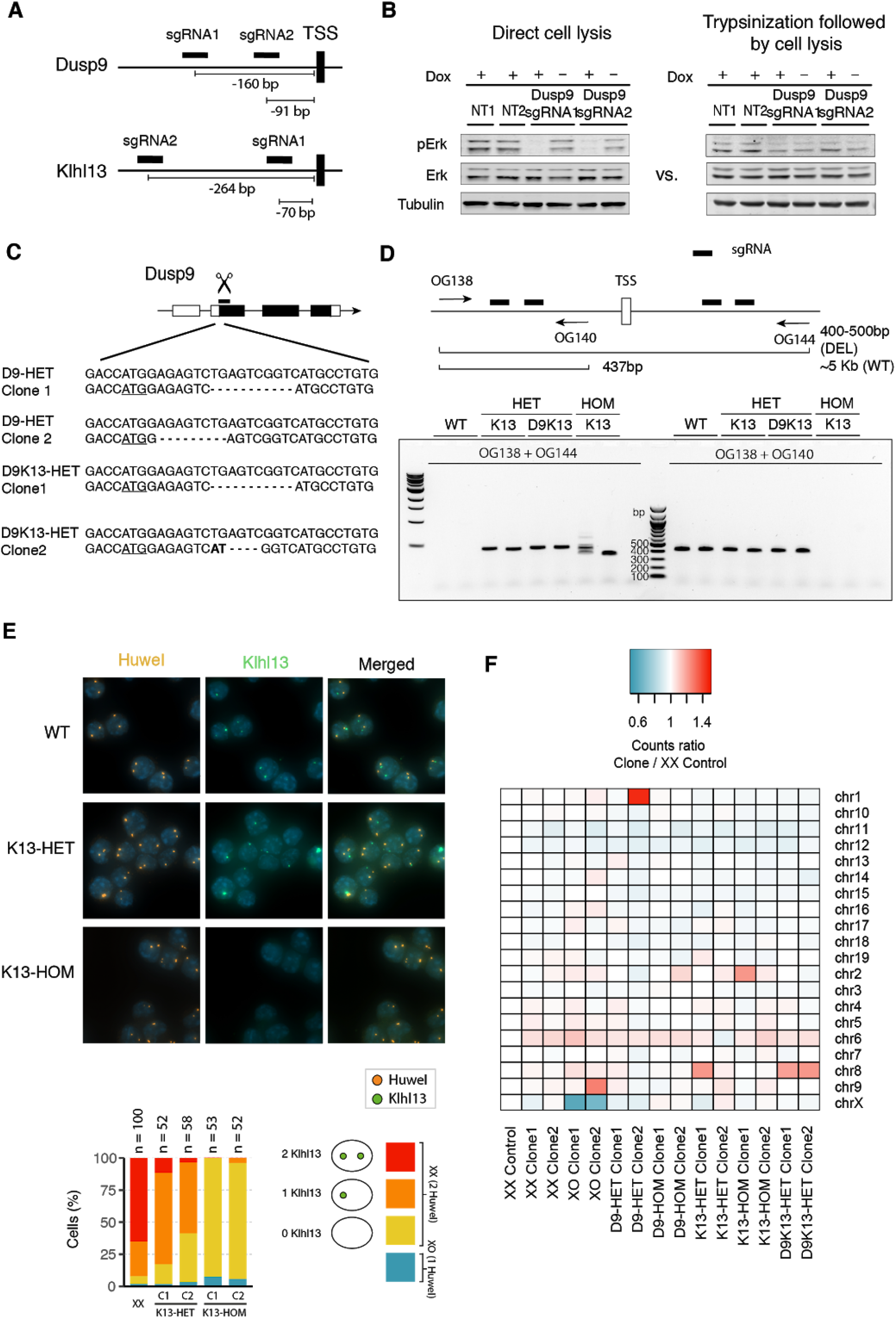
Perturbation of Klhl13 and Dusp9 in mESCs. **(A)** Position of the sgRNA sequences used to over-express Dusp9 and Klhl13 in Fig. 3. SgRNAs for Dusp9 were targeted −91 (chrX:73,639,328-73,639,346, GRCm38/mm10 Assembly) and −160bp (chrX: 73,639,259-73,639,277) bp, whereas for Klhl13 they were targeted −264 (chrX:23,365,328-23,365,347) and −70bp (chrX:23,365,134-23,365,152) bp upstream of the transcriptional start site (TSS) **(B)** Comparison of pErk levels in cells over-expressing Dusp9, harvested either through direct cell lysis (left) or after trypsinization and cell pelleting (right). **(C)** Sequence of Dusp9 frameshift mutants, the start codon is underlined. **(D)** PCR-genotyping of Klhl13 mutant lines. Arrows indicate primer position, thick bars sgRNA target sites. **(E)** RNA fluorescence in situ hybridization of wildtype, heterozygous and homozygous Klhl13 mutant mESCs using probes for Klhl13 (green) and another X-linked gene (Huwe1, orange). The fraction of cells with a specific pattern was quantified (bottom). **(F)** Karyotyping of all cell lines used in Fig. 4 by double digest genotyping by sequencing (ddGBS). Counts mapping to each chromosome were normalized to an XX clone that had previously been karyotyped via metaphase spreads.

**Supplementary Figure 5.**
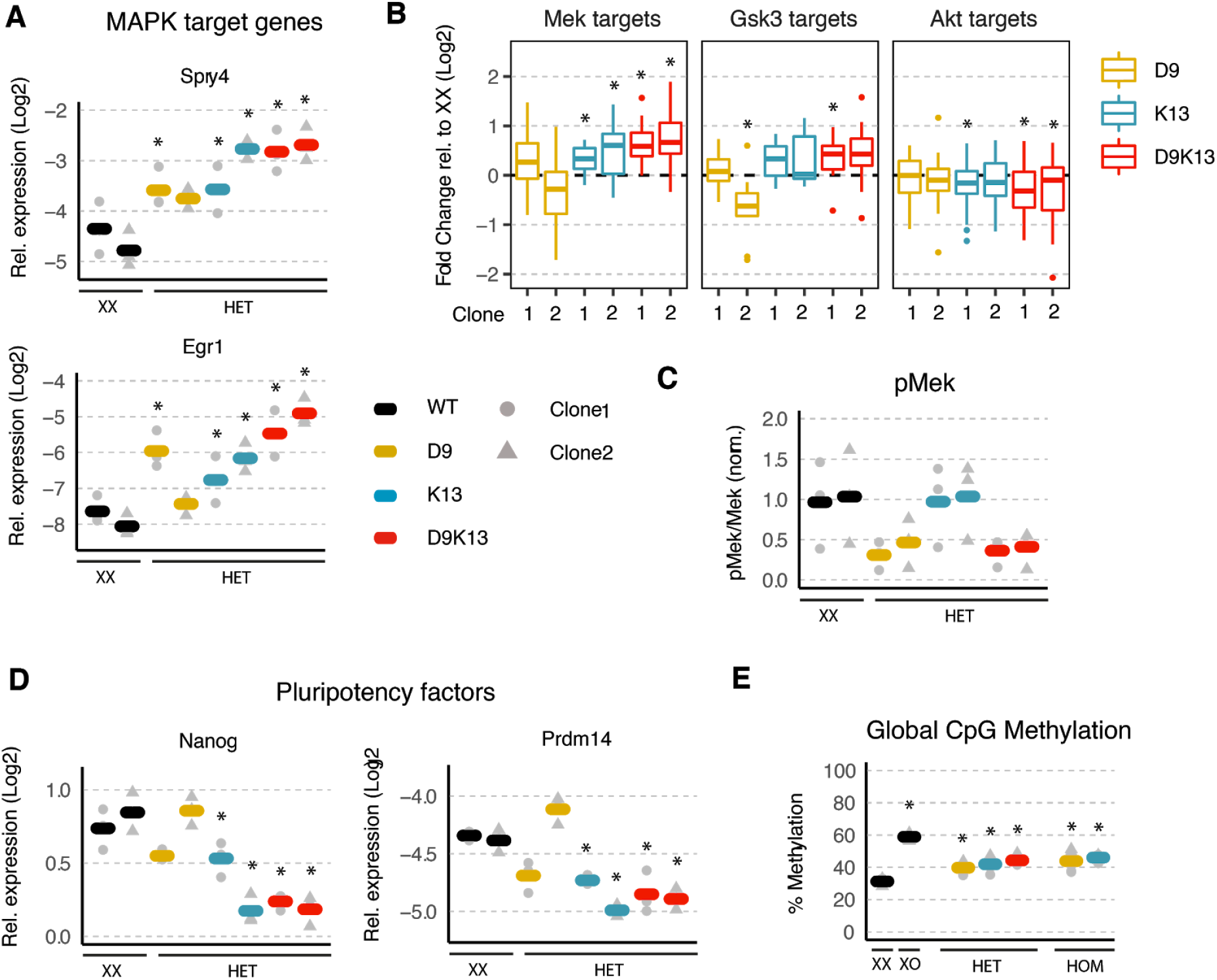
Heterozygous mutations of Klhl13 and Dusp9 in female mESCs partially phenocopy the male pluripotency state. **(A-D)** Comparison of female 1.8 XX mESCs with a heterozygous (HET) deletion of Dusp9 (yellow), Klhl13 (blue) or both (red) with the parental XX line (2 clones per genotype plotted separately). Individual measurements are shown as grey dots (clone 1) and triangles (clone 2) and the mean across three biological replicates for each clone is indicated by a thick bar. **(A)** Quantification of MAPK target genes by qPCR. **(B)** Boxplots showing expression of Mek (left), Gsk3 (middle) and Akt (right) target genes in each clone for cell lines with the indicated genotype, normalized to the average expression in two XX clones, as assessed by RNA-seq. Boxes indicate the 25th to 75th percentiles and the central line represents the median. **(C)** Quantification of pMek, normalized to total Mek and to the XX control cells by immunoblotting. **(D)** Pluripotency factor expression (Nanog and Prdm14) assessed by qPCR. * p < 0.05 Wilcoxon rank-sum test (B), otherwise two-tailed paired Student’s t-test comparing each clone for a given mutant cell line and the mean of the two analyzed XX wildtype control clones. **(E)** Global CpG methylation levels assessed via pyrosequencing-based luminometric DNA methylation assay (LUMA) in the cell lines used in Fig. 4 in Serum/LIF conditions. Individual measurements are shown as grey dots (clone 1) and triangles (clone 2) and the mean across two clones and three biological replicates is indicated by a thick bar. *p < 0.05 two-tailed paired Student’s t-test comparing each mutant/XO cell line with the XX wildtype cells

**Supplementary Figure 6.**
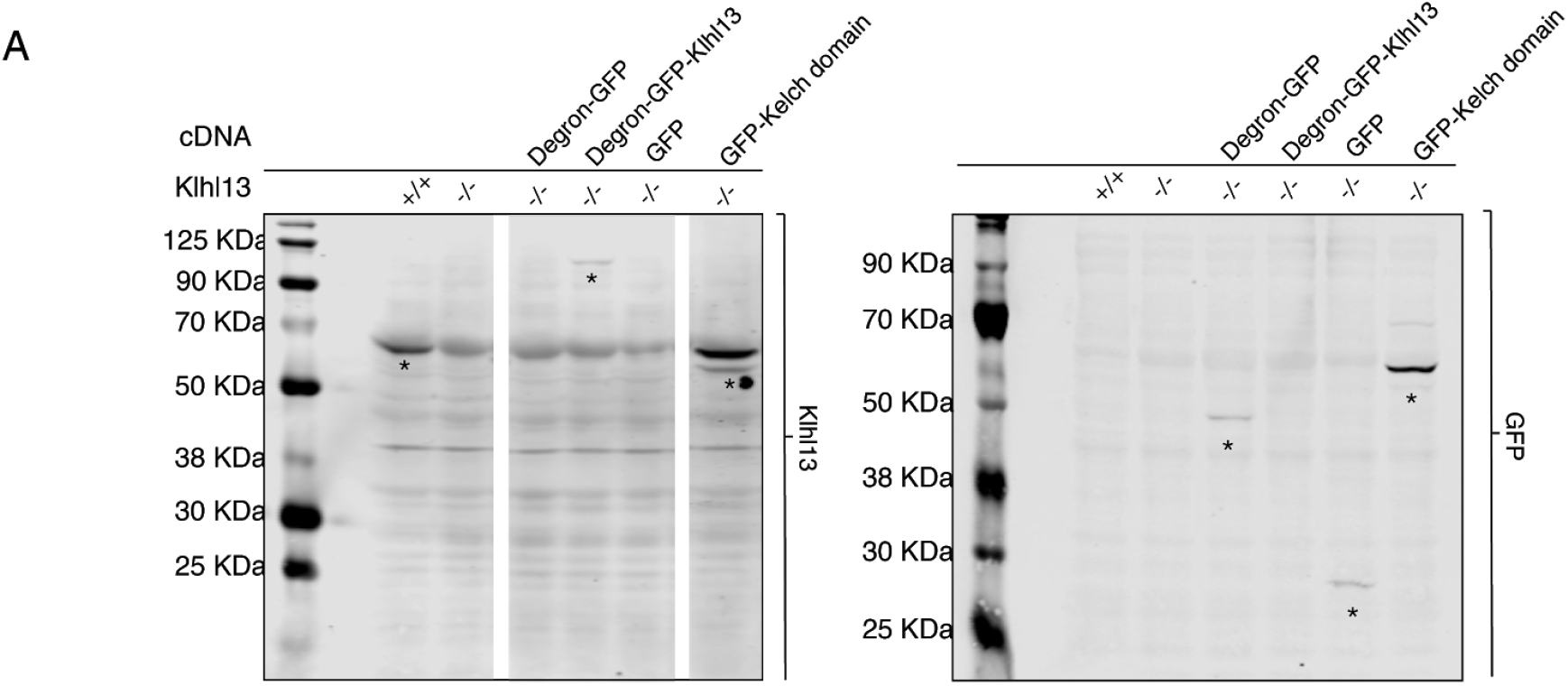
Identification of Klhl13 target proteins that mediate its effect on pluripotency and differentiation. **(A)** Immunoblotting of protein lysates from female K13-HOM cell lines expressing constructs for the identification of Klhl13 interacting proteins and an XX wildtype control. Membranes were incubated with an anti-Klhl13 antibody (left) and an anti-GFP antibody (right). *bands with the expected size.

**Supplementary Figure 7.**
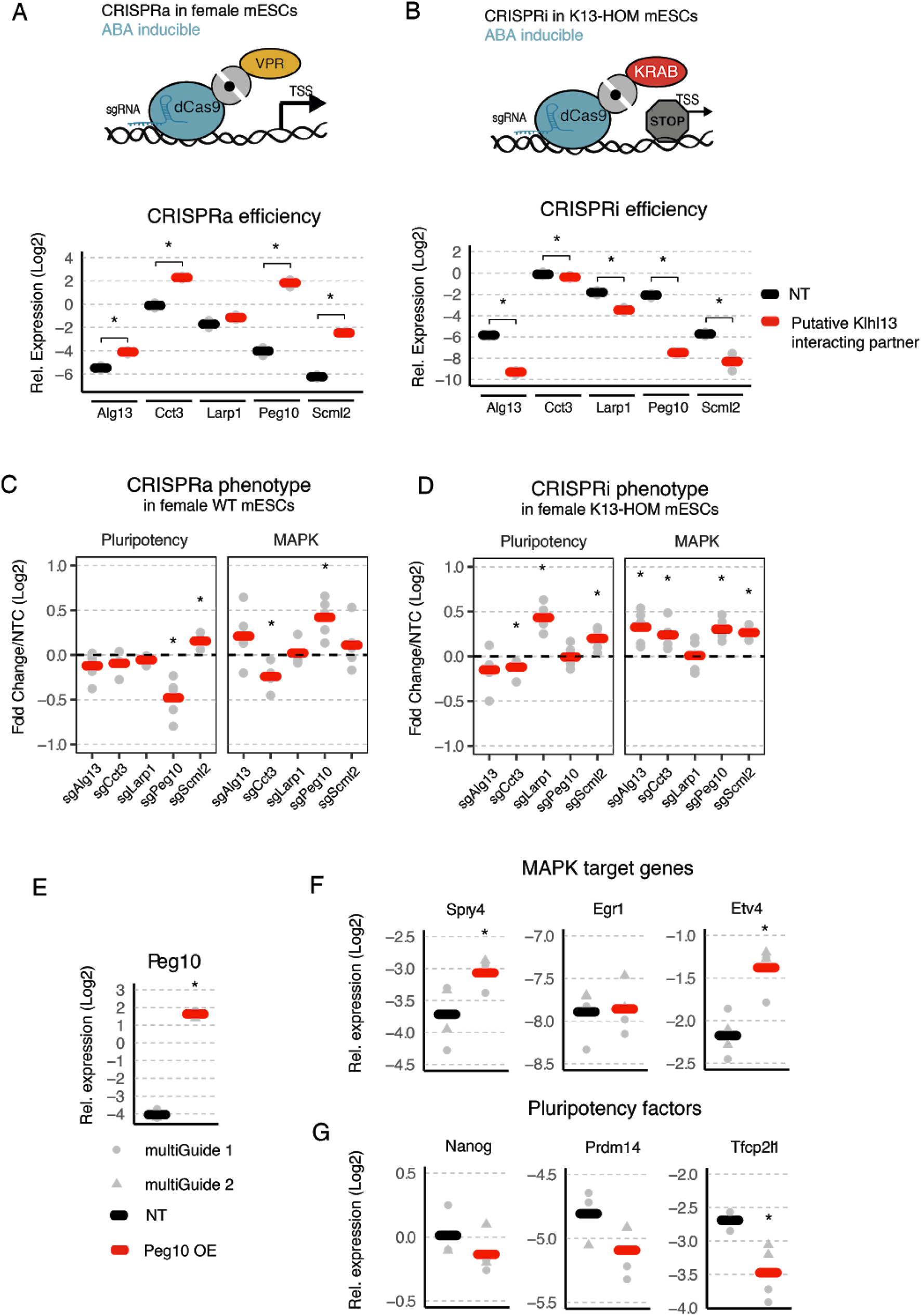
Effects of putative Klhl13 targets proteins on MAPK target gene and pluripotency factor expression. **(A-B)** 1.8 female wildtype mESCs stably expressing the CRISPRa system (A) and 1.8 female K13-HOM mESCs stably expressing the indicated CRISPRi system (B) were transduced with vectors expressing sgRNAs targeting the identified putative Klhl13 target proteins Alg13, Cct3, Larp1, Peg10 and Scml2 (red) or NTCs (black). Expression of each gene was quantified by qPCR in cells expressing the respective sgRNAs or NTCs, as indicated. Individual biological replicates (n=3) are depicted as grey dots and the mean is indicated by a thick bar. Cells were treated with abscisic acid (ABA) for five days prior to cell harvesting for phenotypic assessment. **(C-D)** Expression levels assessed by qPCR of five MAPK target genes (Spry4, Egr1, Etv4, Dnmt3b, Grhl2) and five pluripotency factors (Nanog, Prdm14, Tfcp2l1, Tbx3 and Tcl1) in the cells shown in (A) for (C) and in (B) for (D). Grey dots represent the mean expression of each of the five genes across three biological replicates for cells expressing the respective sgRNAs normalized to the mean expression for cells expressing NTCs for each gene, and bars represent the mean of the five assessed genes. **(E-G)** 1.8 female wildtype mESCs stably expressing the CRISPRa system in (A) were transduced with two different vectors (multiGuide 1 and 2) expressing sgRNAs targeting Peg10 (red) or NTCs (black). Expression levels of Peg10 (E), MAPK target genes (F) and pluripotency factors (G) were quantified by qPCR in cells expressing the two different Peg10 targeting sgRNA or NTC vectors, as indicated. Individual measurements are shown as grey dots (multiGuide construct 1) and triangles (multiGuide construct 2) and the mean across the two multiGuide constructs and two biological replicates is indicated by a thick bar. Cells were treated with abscisic acid (ABA) for five days prior to cell harvesting for phenotypic assessment. * p < 0.05 one-sample t-test (D-E), otherwise two-tailed paired Student’s t-test comparing gene-specific sgRNAs and NTCs are indicated.

**Table S1. Primary and secondary screen data related to Figure 1 and 2 (A-B)** Sequence composition of the GeCKOx (A) and the GeCKOxs (B) sgRNA library. **(C)** Read alignment summary of samples from primary and secondary screens. **(D)** Enrichment, false discovery rate (FDR), rank and hit summary of the generated primary and secondary screens together with the proliferation effects for the SRE-Elk and Nanog screens (Figure S1G, Figure S2H). Results were generated using MAGeCK CRISPR screen analysis tools. Rank results from HitSelect used to generate the GeCKOxs sgRNA library are also shown.

**Table S2. RNA-seq data related to Figure 2 and 4 (A)** Read alignment summary of RNA-seq data of the 1.8 cell line (XX vs XO) related to Figure 2. **(B)** Read alignment summary of RNA-seq data of the generated mutants cell lines related to Figure 4. **(C)** Gene expression values (rpkm) of three biological replicates of the 1.8 cell line (XX vs XO) related to Figure 2. **(D)** Gene expression values (cpm) of three biological replicates and two independent clones of the generated mutant cell lines related to Figure 4.

**Table S3. Proteomics data related to Figure 6 (A)** LFQ (label-free quantification) protein intensities (log2) from three biological replicates of the IP-MS experiment related to Figure 6C (GFP vs GFP-Kelch domain). Data correction analysis was carried out using the Perseus software (v1.6.1.3) **(B)** LFQ protein intensities (log2) from three biological replicates of the IP-MS experiment related to Figure 6D (GFP vs GFP-Klhl13). **(C)** LFQ protein intensities (log2) from three biological replicates and two independent clones of the K13HOM mutant cell line vs wildtype female cells related to Figure 6E. **(D)** Differential expression analysis (Enrichment, p-value and hit summary) from proteomics data related to Figure 6C-E. DE analysis was carried out using the Perseus software (v1.6.1.3)

**Table S4. Reagents and cell lines related to Figure 1-6. (A)** Antibodies used in the study. **(B)** Primer sequences. **(C)** gRNA sequences. **(D)** Cell lines. **(E)** Plasmids. **(F)** Oligo sequences.

